# Changing Pacific salmon nursery lake ecosystem dynamics over centuries to millennia: insights from sedimentary DNA metabarcoding

**DOI:** 10.64898/2026.04.16.718307

**Authors:** Yuanyu Cheng, David Walsh, Joanna Gauthier, Daniel Selbie, Irene Gregory-Eaves

**Affiliations:** Department of Biology, McGill University, Montreal, Quebec, Canada; Groupe de Recherche Interuniversitaire en Limnologie (GRIL), Montreal, Quebec, Canada; Department of Biology, Concordia University, Montreal, Quebec, Canada; Department of Biology, Trent University, Peterborough, Ontario, Canada; Fisheries and Oceans Canada, Science Branch, Pacific Region, Ecosystem Sciences Division, Cultus Lake Salmon Research Laboratory

**Keywords:** Sedimentary DNA, DNA metabarcoding, Pacific Salmon, Paleolimnology, Ecosystem dynamics

## Abstract

Pacific salmon are keystone species to North Pacific freshwater, coastal, and oceanic ecosystems, but many populations have declined or become more variable in recent decades due to anthropogenic impacts and climate change. Long-term records are needed to understand past changes, identify ecosystem stressors, and guide restoration. We used sedimentary DNA (sedDNA), an emerging paleoecological approach offering broader taxonomic information than traditional methods, to reconstruct ecosystem changes across five Pacific salmon nursery lakes in British Columbia (Canada). DNA metabarcoding targeting the 18S ribosomal RNA gene V7 region was used to track shifts in eukaryotic communities including algae and invertebrates over centuries to millennia. Most lakes showed notable algal community shifts over the past two centuries, with declining green algae and rising diatom relative abundances. Chrysophytes and dinoflagellates also increased over the past century in most lakes, likely driven by stronger thermal stratification, which favored these motile and mixotrophic algae that are capable of vertical migration and flexible nutrient acquisition. We contextualized the trajectories of each core through an ordination analysis based on 98 lakes distributed across British Columbia, which identified land-use changes and longer growing seasons as potential drivers. Network analyses of the sedDNA time series revealed decreasing modularity and increasing connection across lakes, suggesting a shift in resilience mechanisms from between-module buffering by compartmentalized specialists to within-guild insurance via functional overlap among generalists. Our findings demonstrate that sedDNA provides taxonomically rich, long-term insights into aquatic ecological dynamics, which are foundational for understanding and protecting Pacific salmon nursery habitats.

## INTRODUCTION

Pacific salmon are an iconic feature of the North Pacific Rim, serving essential ecological, cultural, and economic roles (Schindler et al. 2003). Ecologically, these anadromous species support remarkable biodiversity across their range. They contribute to marine ecosystems where they mature, to freshwater systems where they spawn, and to riparian forests enriched by their carcasses. They function as vital nutrient vectors that transfer marine-derived resources to terrestrial and freshwater food webs (Gende et al. 2002). Culturally, Pacific salmon have sustained Indigenous peoples of the Pacific Northwest for millennia, forming the foundation of traditional diets, ceremonies, and economies, and remaining central to Indigenous identity and sovereignty (Moss and Cannon 2011). Economically, commercial fishing of Pacific salmon generates hundreds of millions of dollars annually and supports thousands of jobs (Fisheries and Oceans Canada 2024).

The productivity and resilience of salmon populations emerge from hierarchically nested drivers spanning multiple scales, from ocean-basin climate oscillations to regional watershed characteristics and to local nursery lake habitat quality (Mantua 2015; Jones et al. 2020; Price et al. 2024). Accelerating human impacts, including overfishing, habitat disturbance, fish farming, and climate warming, have affected salmon throughout their life cycle, resulting in population declines, reduced body size, and elevated parasitic infection rates in some regions (Krkosek et al. 2007; Oke et al. 2020; Crozier et al. 2021; Ford et al. 2025). Nonetheless, population dynamics of sockeye salmon vary considerably geographically, in part because climate change at the northern end of their range is enhancing productivity but having a negative effect at the southern range limit (Connors et al. 2020).

Effective conservation strategies for Pacific salmon need to account for the complex array of factors affecting populations across freshwater and marine environments. Traditional monitoring approaches, while valuable, face significant limitations: systematic salmon data rarely extends before the 1920s, leaving critical gaps in our understanding of baseline conditions and long-term ecosystem dynamics. Even more limited are data on nursery lake habitat attributes, particularly food-web structure and productivity, which are the ecological foundation supporting juvenile salmon growth and survival. Understanding these food webs is crucial because prey availability and quality, predation pressure, and competitive interactions during freshwater residence directly shape juvenile salmon body size, marine survival, and ultimately recruitment success (Henderson and Cass 1991).

The financial and logistical constraints of long-term monitoring have spurred growing paleoecological research on salmon nursery lakes, where researchers apply diverse biological and geochemical proxies to reconstruct historical salmon populations and their ecosystems (Finney et al. 2002; Selbie et al. 2007). While δ^15^N signatures have successfully reconstructed salmon-derived nutrients and biomass in some pristine systems, this proxy becomes unreliable in lakes with confounding nitrogen sources from human waste inputs or where precipitation-driven terrestrial nitrogen influx varies substantially (Selbie et al. 2009). Similarly, traditional paleoecological approaches to food web reconstruction have relied primarily on preserved subfossils of diatoms and cladocerans (Barouillet et al. 2024), yet many ecologically important algal and zooplankton species or groups leave few to no subfossil remains (e.g., green algae, dinoflagellates, copepods, and rotifers), creating an incomplete picture of ecosystem dynamics (Gregory-Eaves and Smol 2024).

Recent advances in sedimentary DNA (sedDNA) analysis offer unprecedented opportunities to capture this “hidden” biodiversity and reconstruct more comprehensive food web structures (Capo et al. 2021). Among sedDNA approaches, metabarcoding provides a cost-effective method for detecting shifts in community composition with relatively fine taxonomic resolution (Taberlet et al. 2012). Depending on primer selection, metabarcoding can target specific taxonomic groups at species or genus level, or employ universal primers to survey broad assemblages across multiple trophic levels (Domaizon et al. 2017). By applying such molecular techniques, researchers can now reconstruct ecological dynamics of diverse organisms across trophic levels and provide the long-term perspective essential for understanding how food web alterations influence salmon populations under accelerating environmental change (Balint et al. 2018).

We applied DNA metabarcoding targeting the 18S ribosomal RNA gene V7 region to reconstruct changes in eukaryote assemblages across five salmon nursery lakes in British Columbia (Canada) over the past several hundred to thousand years. This marker has proven effective in previous studies, accurately preserving phytoplankton dynamics from monitoring data in sedDNA records and capturing organisms lacking subfossil remains to reveal historical ecosystem dynamics (Capo et al. 2016; Gauthier et al. 2021; Garner et al. 2025). We addressed four research questions: (1) How have algal and zooplankton communities - the primary food web supporting planktivorous juvenile salmon - changed through time? (2) Can fish DNA be detected using this primer set, and what temporal patterns emerge? (3) What environmental drivers explain variation in community dynamics, as inferred by projecting our lakes onto ordination space constructed from 98 British Columbia lakes with contemporary data? (4) How did functional groups shift over time and change network properties (e.g., modularity, connectance)? Through these reconstructions of long-term ecosystem dynamics, we identified centennial- to millennial-scale trends and potential drivers that have shaped these critical habitats, providing essential baseline data for understanding contemporary changes and informing conservation strategies for salmon nursery lakes.

## MATERIALS AND METHODS

### Study lakes

Our study lakes are in British Columbia (BC), Canada (Fig. 1a), spanning a latitudinal gradient, from Cultus Lake in the South and Tahltan Lake in the North. These lakes exhibit diverse morphologies, watershed characteristics, and limnological properties (Table 1). Babine and Shuswap lakes are large and deep; Babine is the longest natural lake in BC with a length of 153km, while the remaining three lakes are smaller with moderate depths (Table 1). This north-south distribution captures a gradient of climate contexts and anthropogenic influences (Fig.1b).

**Fig. 1.**
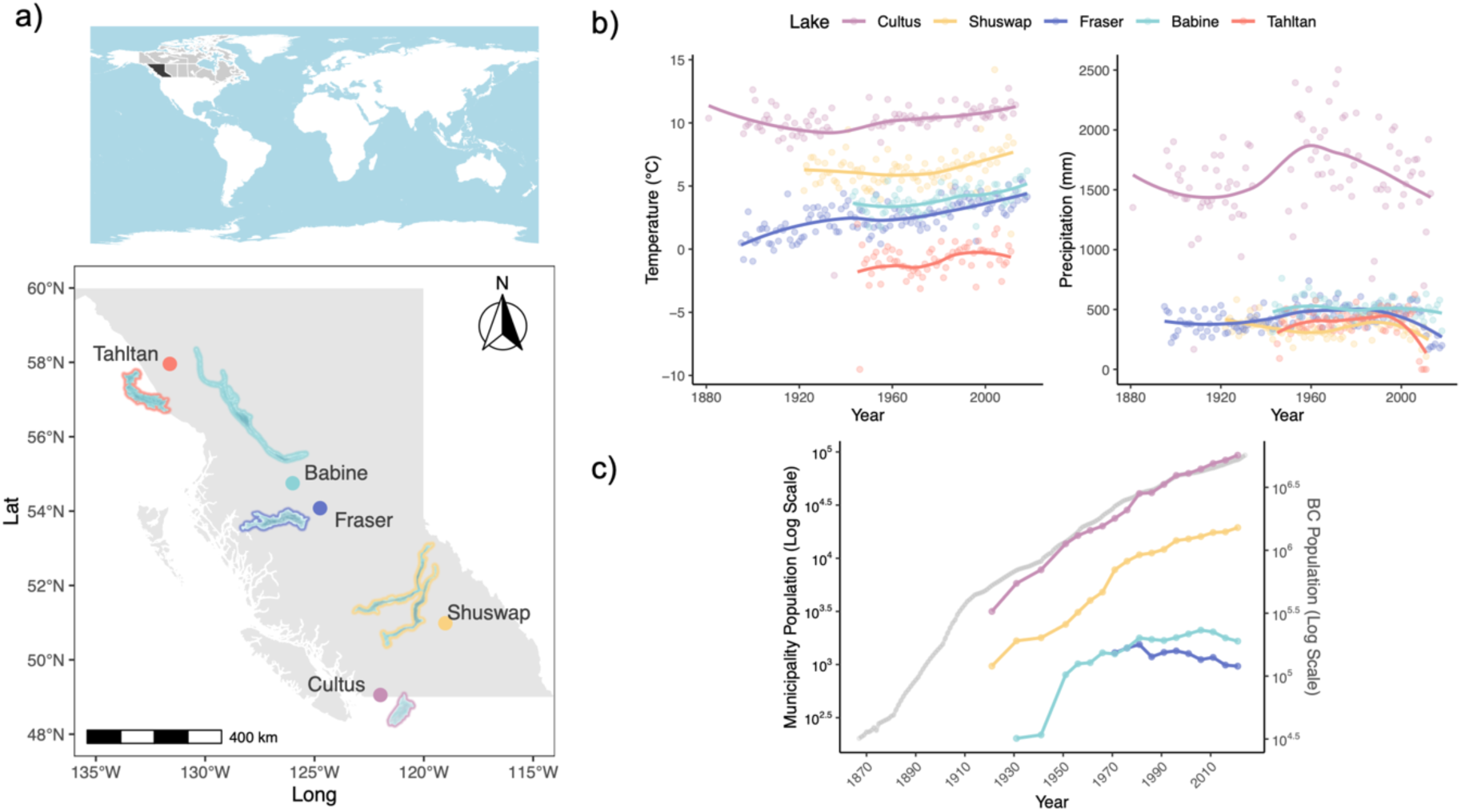
Locations of the study region in British Columbia (BC), Canada (top panel), and the study lakes (reference panel a, lakes not drawn to scale). b) Long-term climate (air temperature and precipitation) in the study region. Annual averages (dots) with loess smoothed trend line. c) Human population trends for the province of BC in grey and the municipalities adjacent to each study lake (Tahltan Lake region sparsely populated; data not available).

**Table 1.** Limnological characteristics of the five study lakes. Water chemistry data were obtained from the BC monitoring network and are presented as mean values (± SD) across hypolimnion and epilimnion samples collected in spring and summer over multiple years. Average sockeye spawner counts (10-year means) were obtained from the Salmon Escapement Database and standardized by lake area (km²) to calculate spawner density.

| Lake | Lat | Long | Elevation<br>(m) | Watershed | Max<br>depth<br>(m) | Area<br>(km<sup>2</sup>) | TP<br>(μg/L) | DOC<br>(mg/L) | Average<br>spawners<br>count (#) | Spawner<br>density<br>(#/km<sup>2</sup>) |
| --- | --- | --- | --- | --- | --- | --- | --- | --- | --- | --- |
| Cultus | 49.0533 | -121.9867 | 47 | Fraser | 42 | 6 | 4.5<br>(± 1.5) | 1.4<br>(± 0.3) | 1,092 | 173 |
| Shuswap | 50.9833 | -119.0167 | 347 | Fraser | 265 | 310 | 4.2<br>(± 1.1) | 2.2<br>(± 0.3) | 505,669 | 1,633 |
| Fraser | 54.0784 | -124.7465 | 670 | Fraser | 30.5 | 54 | 19.4<br>(± 7.2) | 8.4<br>(± 0.6) | 134,818 | 2,497 |
| Babine | 54.7500 | -126.0000 | 711 | Skeena | 186 | 479 | 3.8<br>(± 0.9) | 8.0<br>(± 0.4) | 73,050 | 153 |
| Tahltan | 57.9588 | -131.6189 | 811 | Stikine | 48 | 5 | 10.6<br>(± 1.8) | | 27,435 | 5,599 |

Long-term climate records were obtained from the nearest climate station of each lake with continuous data spanning at least 70 years (station information in Table S1). Regional trends show modest increases in annual air temperature and decreases in precipitation over time (Fig. 1b). Mean annual temperature follows a latitudinal gradient, with Cultus (southernmost lake) exhibiting the highest values and Tahltan (northernmost) the lowest. Precipitation levels were historically similar across regions except for Cultus, which receives substantially more precipitation, predominantly as rain (∼6% as snow vs. 24-49% for other lakes). All lakes have experienced declines in precipitation over recent decades, with the most pronounced rate of change in precipitation since the ∼2000s apparent at Tahltan (Fig. 1b).

The north-south distribution of our study lakes captures a gradient of anthropogenic influences. Southern systems (Cultus and Shuswap lakes) have experienced rapidly increasing human populations over the past century, while northern systems (Fraser and Babine lakes) have maintained consistently lower population densities, with recent declines (Fig. 1c). The Tahltan Lake region is likely the least populated due to its remote location, though long-term census data are unavailable. Cultus Lake, within the territory of Soowahlie First Nation and the Fraser Valley Regional District, supports intensive recreation in a region with farming history (details in Gauthier et al. 2021) and has experienced substantial increase in nutrient loadings (Putt et al. 2019). Shuswap Lake, named after the Shuswap Indigenous Peoples (the Secwepemc), supports extensive recreation, lakeside communities, and modest agriculture. Fraser Lake, in the Nadleh Whut’en territory, is near to a village of less than 1,000 people. Nonetheless, Fraser Lake has been undergoing cultural eutrophication over the past few decades, likely driven by agricultural and residential runoff (Hobbs and Wolfe 2008). Babine Lake lies within the traditional territory of Lake Babine Nation and has dispersed and small human settlements. Tahltan Lake, within the Tahltan Nation territory, represents the most pristine site, remote from major urban centres.

Fisheries enhancement efforts have varied across sites, with Babine subjected to major spawning channel construction and fry stocking from the late 1960s onwards (Babine Lake Development Project, BLDP; Cox-Rogers and Spilsted 2012), while Tahltan and Cultus experienced fry stocking programs (Fig. S1). Additional local history details are provided in Supplementary Information S1 and Fig. S1.

### Sediment sampling

Sediment cores were collected from depositional basins within each of the study lakes in fall 2023 (Babine, Cultus, Fraser, and Shuswap Lakes) and in summer 2024 (Tahltan Lake), using a Super Glew gravity corer (internal diameter: 8.6 cm). Sediment core lengths ranged from ∼30 to 45 cm. Zorbitrol was applied to the sediment-water interface in each core to preserve stratigraphic integrity. All sediment cores were sealed in black plastic bags upon recovery to limit light exposure, and packed in a styrofoam cooler with chiller packs followed by rapid shipment to McGill University, where they were stored at 4°C until core splitting and subsampling.

Subsampling was conducted following strict sedimentary ancient DNA (sedaDNA) protocols (Garner et al. 2025). Prior to core splitting and subsampling, all lab surfaces were sterilized with bleach (1%), triple-rinsed with deionized water, and then sanitized using DNAaway (Thermo Fisher). Each sediment core was split, with one half archived and the other designated for molecular and geochemical analyses. For molecular analyses, sediments were immediately sub-sampled by collecting the innermost portion with a sterile spatula, transferred to sterile 50 mL tubes, and stored at −20 °C. The remaining sediments of each interval were placed in Whirl-Pak bags, freeze-dried, and preserved for geochemical analyses.

### Geochemical analyses

Sediment chronologies were established upon the radioisotope activities of ^210^Pb (lead), ^137^Cs (Cesium), ^214^Pb, and ^214^Bi (Bismuth), measured by gamma spectrometry (ORTEC-GWL high-purity germanium photon well detector with HPLB shielding, coupled to a Digital Gamma-Ray Spectrometer; DSPEC Jr. 2.0) at Fisheries and Oceans Canada Cultus Lake Salmon Research Laboratory. ^210^Pb activities generally declined exponentially with depth across all cores, with minor irregularities indicating variations in sedimentation rates (Fig. S2). ^210^Pb ages were determined using the Constant Rate of Supply (CRS) model (Fig. S2; Appleby 2001). For Cultus, Fraser, and Tahltan, polynomial models based on ^210^Pb ages were used to extrapolate the ages of deeper samples without CRS dates (Fig. S3); we interpreted the extrapolated ages with caution. For Tahltan, the polynomial model was constrained by one ¹⁴C date from pollen extracts prepared by the LacCore facility at the University of Minnesota (details in Fig. S3). For Babine, insufficient data prevented extrapolation beyond ∼1700 CE; older intervals are presented as pre-1700 CE. For Shuswap, ²¹⁰Pb dating covered the entire analyzed sediment core length, eliminating the need for polynomial extrapolation.

### Molecular analyses

DNA extraction was performed using the Qiagen PowerMax kit, following the manufacturer protocol in a dedicated clean room. Extraction blanks were run every 19th sample alongside the sediment samples. The eluted DNA was concentrated using Amicon Ultra-15 centrifugal filter units at 4000 × g for 5 minutes at room temperature to a final volume of ∼100-200ul. Downstream PCR amplification, library preparation, and Illumina sequencing were carried out at the McGill Genome Centre.

The V7 region of the 18S rRNA gene was amplified using the primer set 960F (5’-GGCTTAATTTGACTCAACRCG-3’; Gast et al. 2004) and NSR1438 (5’-GGGCATCACAGACCTGTTAT-3’; Van de Peer et al. 2000). Each 25 µL PCR reaction contained 3 µL of DNA template, 1× Qiagen PCR buffer, 0.2 mM dNTP mix, 0.2 µM of each primer, 2.5 U Taq DNA polymerase, 2.5 mM MgCl₂, and 1× Qiagen Q-solution (for highly inhibitory samples), with nuclease-free water added to reach the final volume. PCR cycling conditions were: initial denaturation at 94°C for 3 minutes; 30 cycles of 94°C for 30 seconds, 55°C for 1 minute, and 72°C for 1 minute; followed by final elongation at 72°C for 10 minutes.

Library preparation was performed in a total reaction volume of 25 µL. Each reaction consisted of 2.5 µL of the first PCR product diluted 1:20, 2 µM of index forward primer, 2 µM of index reverse primer, 12.5 µL of 2× Kapa HiFi HotStart ReadyMix, and 5 µL of nuclease-free water. Cycling conditions were: initial denaturation at 95°C for 3 minutes; 12 cycles of 95°C for 30 seconds, 55°C for 30 seconds, and 72°C for 30 seconds; followed by final elongation at 72°C for 5 minutes. Sequencing was performed on an Illumina MiSeq platform using 2 × 250 bp paired-end reads. Extraction blanks were sequenced to assess contamination.

### Bioinformatic analyses

The demultiplexed raw sequence data were processed using cutadapt version 5.1 to trim primers (Martin 2011). The DADA2 package version 1.30.0 (Callahan et al. 2016) in R software (version 4.3.0, R Core Team 2023) was then used to filter reads, merge paired end sequences, and remove chimeras. Sequences with fewer than 10 total reads were removed to reduce sequencing artifacts or spurious reads. Taxonomic identification was performed using the MOTHUR classify.seqs algorithm with the Protist Ribosomal Reference (PR2 version 5.0.0) SSU rRNA gene database (Guillou et al. 2013), applying a minimum bootstrap confidence threshold of 80%. For higher taxonomic resolution, BLASTn top hit analysis was employed. When multiple hits yielded identical bitscores, we constructed a phylogenetic tree of the top ASVs within that taxonomic group and assessed congruency among the top hits to assign taxonomy. Only the top 1,000 ASVs ranked by abundance across the entire dataset were included in BLASTn top hit analysis.

### Statistical analyses

All statistical analyses were performed in R version 4.3.0 (R Core Team 2023). Community composition was visualized using heat tree diagrams generated with the metacoder package (Foster et al. 2017). Temporal trends across study lakes were examined through stacked bar plots at multiple taxonomic levels (e.g., algae, zooplankton, fish) using ggplot2 version 3.5.2 (Wickham 2016).

Non-metric multidimensional scaling (NMDS) ordination was conducted using our salmon lake biological dataset previously Hellinger-transformed, to evaluate temporal changes in community composition, and assess the distribution of dominant taxonomic classes with ecological implications (vegan package version 2.7-1, Oksanen et al. 2020). To investigate potential environmental drivers of these observed patterns, we employed a passive plotting approach, using the NSERC Canadian Lake Pulse Network (hereafter LakePulse) water column dataset, that applied 18S rRNA metabarcoding using the same primer set (Garner et al. 2022).

LakePulse was a pan-Canadian survey that sampled hundreds of lakes once during summer 2017-2019, and 18S metabarcoding was conducted on DNA extracted from filtered water samples with the same primer set used in our study (details in Garner et al. 2022). Briefly, water samples were collected using an integrated tube sampler from the euphotic zone over a depth of up to 2m below the surface at the deepest point of each lake. Water samples were prefiltered through 100-µm nylon mesh, then vacuum filtered on 47-mm-diameter 0.22-µm Durapore membrane,s and immediately put in a −80°C freezer until processing. Of our study lakes, only Cultus Lake was part of the lakes sampled in the LakePulse Network.

We restricted our analyses to the LakePulse datasets within the province of British Columbia (*n* = 98). Overall, the integration of such large datasets addresses a common limitation in paleolimnological studies, where environmental data are often lacking for interpreting long-term sediment core records. We created an ordination using LakePulse eDNA data (Hellinger-transformed) as the framework, then passively projected both LakePulse environmental variables (water chemistry and land-use data) and our 18S rRNA data from the five lakes. To align the two 18S rRNA datasets, we filtered our sedDNA dataset to retain only amplicon sequence variants (ASVs) that matched those in the LakePulse project. Environmental variables from the LakePulse dataset were transformed as appropriate (e.g., log, natural log, square root) to best approximate normal distributions, and then screened for significance using the envfit() function in vegan R package (Oksanen et al. 2020). Variables with permutation test p-values less or equal to 0.05 were considered statistically significant. Downcore salmon site scores were passively projected into the ordination space using weighted averaging based on matched ASV abundances (Hellinger-transformed).

Stratigraphically constrained cluster analysis (CONISS) with broken-stick model validation was conducted to delineate major ecological zones using the rioja package version 1.0-7 (Juggins 2022). When the broken-stick model did not indicate statistically significant divisions, zones were determined through visual inspection of the dendrogram. Identified time zones for each lake are shown in Fig. S4.

Network analyses were conducted using the igraph (version 2.1.4; Csárdi et al. 2025) and ggClusterNet2 (version 0.1.0; Wen et al. 2025) packages to investigate taxa co-occurrence patterns and potential interactions across the identified time zones with CONISS. To minimize bias from variable sequencing depths, all samples were rarefied to 36,146 reads per sample, excluding five older samples from Tahltan and Babine with low read abundances. Within each time zone, the 100 most abundant ASVs were selected for network construction. Spearman rank correlations were calculated between ASVs, and statistical significance was assessed through permutation testing with 100 random network samplings and 10 repetitions. Correlations with coefficients |r| > 0.6 and p-values < 0.05 were retained for network construction. Network communities were detected using the fast greedy modularity optimization algorithm (Wen et al. 2025), and networks were visualized with functional group assignments from Garner et al. (2022). Topological roles (connectors, module hubs, network hubs, and peripherals) of each network were identified using the ggClusterNet2 package.

## RESULTS

### 18S rRNA gene sequence statistics

After bioinformatic processing, 9,298,026 sequences corresponding to 11,241 amplicon sequence variants (ASVs) were obtained from the sediment samples. In the blank controls, 435 sequences (< 0.005% of total sequences) from 7 ASVs were detected (Fig. S5). Following a correction, whereby the maximum abundance of each ASV detected in the blank controls were subtracted from all sediment DNA samples, 9,289,496 sequences remained (99.91% of the original dataset).

### Taxonomic diversity based on the 18S rRNA gene metabarcoding

We detected substantial taxonomic diversity across the five study lakes spanning multiple major eukaryotic lineages (Fig. S6), with diverse and abundant representation of Alveolata (1325 ASVs, 17% of sequences), Rhizaria (1260 ASVs, 6% of sequences), Fungi (1247 ASVs, 8% of sequences), Gyrista (988 ASVs, 20% of sequences), and Metazoa (495 ASVs, 25% of sequences). Within Alveolata, we observed a diverse assemblage of ciliates alongside dinoflagellates and parasitic Apicomplexa. Rhizaria showed limited taxonomic resolution, with broad categories of Cercozoa and Foraminifera represented. Gyrista contained several major lineages, including diatoms (Bacillariophyceae, Coscinodiscophyceae, and Mediophyceae), golden algae (Chrysophyceae), algal parasitoids (Pirsoniales), and other taxa. Green algae were well-represented within Chlorophyta, while the closely related Streptophyta included various plant groups. Amoebozoa was represented by multiple clades, including Discosea, Evosea, and Tubulinea. Finally, Opisthokonta was dominated by metazoan and fungal taxa, with smaller contributions from Ichthyosporea and choanoflagellates.

When data from all lakes were combined, Arthropoda (173 ASVs, 20% of sequences) emerged as the most abundant class, followed by algae, microfungi, and parasites (including algae-affecting Pirsoniales and arthropod-affecting Gregarinomorphea). However, individual lake patterns revealed considerable variation from this overall trend (Fig. S7). The southern lakes (Cultus and Shuswap) were characterized by high Arthropoda dominance, while the northern lakes (Fraser, Babine, and Tahltan) displayed more balanced distributions among taxonomic classes, with algae predominating (Fig. S7). Within the northern lakes, the dominant algal groups differed markedly: Chrysophyceae dominated in Fraser Lake, Coscinodiscophyceae in Babine Lake, and Dinophyceae in Tahltan Lake (Fig. S7).

### sedDNA-inferred temporal trends in phytoplankton

Unicellular algae are abundant in pelagic food webs and serve as foundational conduits of energy to higher trophic levels (e.g., zooplankton and fish), thus contributing substantially to the sedDNA pool captured in our profundal cores. Here, we focused on the major algal taxa: Bacillariophyta (diatoms), Chlorophyta (green algae), Chrysophyta (golden algae), Cryptophyta (cryptophytes), and Dinophyta (dinoflagellates). Cryptophytes maintained consistently low detection levels across all systems, while other taxa exhibited discernible trends (Fig. 2a).

**Fig. 2.**
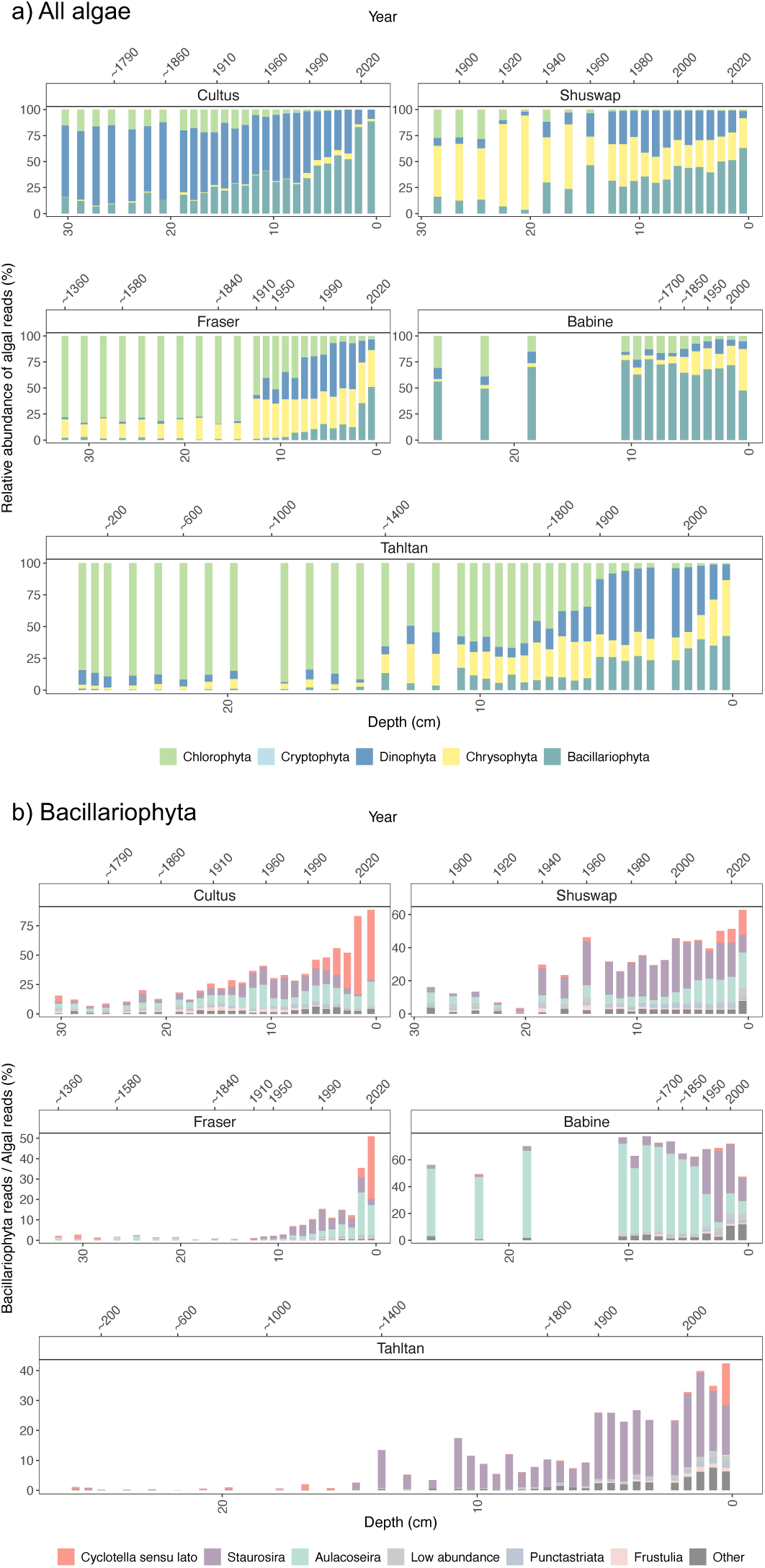
Bar charts showing temporal changes in: a) the relative abundance of the five major algal groups (Chlorophyta, Cryptophyta, Dinophyta, Chrysophyta, Bacillariophyta) and b) Bacillariophyta genera as a proportion of total algal reads. ASVs beyond the top 1000 are grouped as low abundance.

Diatoms increased over time across most lakes, in terms of relative sequence abundance, with the notable exception of Babine Lake, which has been consistently dominated by diatoms throughout the past millennium (Fig. 2). In the three more southern lakes, consistent two-phase pattern emerged: an initial increase in both the benthic taxon *Staurosira* and the tychoplanktonic genus *Aulacoseira*, followed by a recent increase in planktonic diatoms (i.e., *Cyclotella* sensu lato; shown in red; Fig. 2b), particularly over the past two decades. Due to high genetic similarity in the 18S rRNA gene v7 region, genera within the *Cyclotella* sensu lato (e.g., *Stephanodiscus*, *Lindavia*) cannot be reliably differentiated and are grouped together, despite their distinct nutrient autecology (Cumming et al. 2015). Babine Lake, by contrast, yielded a sedDNA diatom record dominated by *Aulacoseira* throughout, with a recent increase in *Staurosira*. In the northernmost Tahltan Lake, *Staurosira* increases in relative abundances over the past ∼500 years and in the past two decades we see an increase in the *Cyclotella* sensu lato.

Chlorophytes exhibited declining trends across all systems (Fig. 2a). This pattern was most pronounced in Fraser and Tahltan lakes, where chlorophytes historically comprised over 70-90% of total algal reads but declined to below 3% in recent decades (Fig. 2a). The dominant genera within this group were *Desmodesmus* and *Choricystis*.

Chrysophytes exhibited an overall recent increasing temporal trend across most lakes, of varying sedDNA relative read abundances (Fig. 3a). Chrysophytes remained scarce in Cultus Lake (<7% of total algal reads in any interval), while being more abundant in other lakes (>30-60% in certain intervals; Fig. 3a). The chrysophyte communities based on the sedDNA analyses shifted from dominance by a single genus (*Ochromonas*) to a more diverse assemblage including *Ochromonas*, *Mallomonas*, *Spumella*, *Dinobryon*, and *Chrysosphaera* after the 1950s in most lakes (Fig. 3a).

**Fig. 3.**
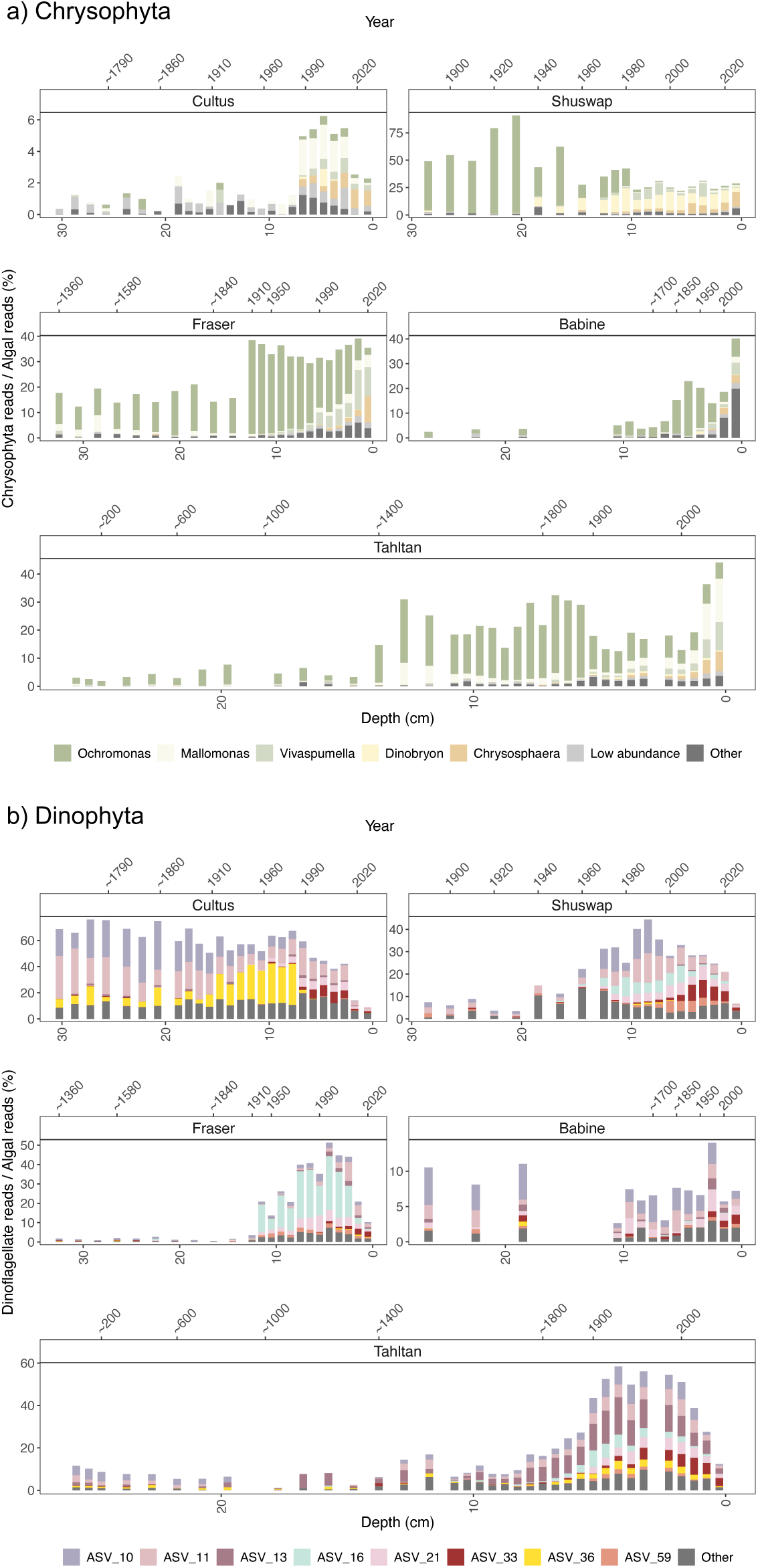
Bar charts showing temporal changes in: a) Chrysophyta genera, and b) Dinophyta ASVs as a proportion of total algal reads. ASVs beyond the top 1000 are grouped as low abundance.

Dinoflagellates showed divergent lake-specific trends over the past two centuries (Fig. 3b). They dominated Cultus Lake until the late 20th century, while in Babine Lake they remained consistently low in abundance, generally below 15% of total algal reads. In Shuswap, Fraser, and Tahltan lakes, dinoflagellates exhibited initial increases beginning in the early-to-mid 20th century, followed by recent declines. Taxonomic identification at the genus level was challenging due to genetic similarity among dinoflagellates with the marker used; however, analysis at the ASV level revealed distinct changes in community composition across lakes and through time. For example, ASV33 (in dark red) increased in relative read abundance over the past three decades across all lakes (Fig. 3b).

### Temporal trends in other taxonomic groups

Arthropoda accounted for substantial sequence reads across lakes (Fig. S7), with Copepoda being the most dominant taxon (Fig. S8). Copepods dominated sedDNA-inferred zooplankton communities in Cultus Lake (average 47% of total sequences) and Shuswap Lake (average 21% of total sequences). In both of these southern lakes, *Epischura* copepods were the dominant taxon, represented primarily by two genotypes (ASV1 and ASV7). Fraser Lake exhibited a similar copepod assemblage, but at much lower relative abundance (average 8% of total sequences), while Babine Lake exhibited only sporadic, low-abundance copepod detections (average 2% of total sequences). In contrast, Tahltan Lake supported a distinct copepod assemblage dominated by *Leptodiaptomus*, with peaks occurring during two distinct periods: around 1200-1400 CE and 1700-1900 CE. Across all lakes, rotifers contributed far fewer sequence reads than copepods (84,863 vs. 1,753,724 total sequences, respectively; Fig. S8). Cladocerans (*Holopedium* and *Bosmina*) were detected infrequently across all lakes, totaling only 5 ASVs corresponding to 5,493 sequences across all study sites (Fig. S8).

Fish sequences were relatively low in abundance, comprising 10 ASVs and a total of 6,368 reads, but exhibited varying patterns in relative read abundance across the nursery lakes (Fig. S9). The dominant species detected was sockeye salmon (*Oncorhynchus nerka*), represented by 4 ASVs and 4,908 reads. Other detected species included shorthead sculpin (*Cottus confusus*), coho salmon (*Oncorhynchus kisutch*), goby (*Eucyclogobius newberryi*), plains sucker (*Pantosteus jordani*), and cutthroat trout (*Oncorhynchus clarkii lewisi*). In general, fish relative read abundance remained low overall but was slightly higher historically across most lakes, except at Babine Lake where recent intervals showed elevated fish sedDNA reads (Fig. S9).

### Temporal shifts in eukaryotic communities

Non-Metric Multidimensional Scaling (NMDS) ordination reveals consistent directional temporal trends across all five lakes, with trajectories pointing toward the bottom left quadrant despite different starting positions (Fig. 4a). To identify how the eukaryotic assemblage changed over time, we plotted the amplicons with relatively high abundance (i.e., > 4% in at least 2 samples) and high loadings on the NMDS axes (i.e., > 0.5; Fig. 4b). With this analysis, we can see that the older intervals of the Babine Lake sediment core were characterized by *Aulacoseira granulata* (ASV32, ASV47, and ASV64), whereas more recent samples were represented by *Aulacoseira subarctica* (ASV23), a taxon that also appeared in recent samples from other lakes (Fig. 4b). *Cyclotella* sensu lato (ASV18) showed increasing trends across all lakes over time (Fig. 4b).

**Fig. 4.**
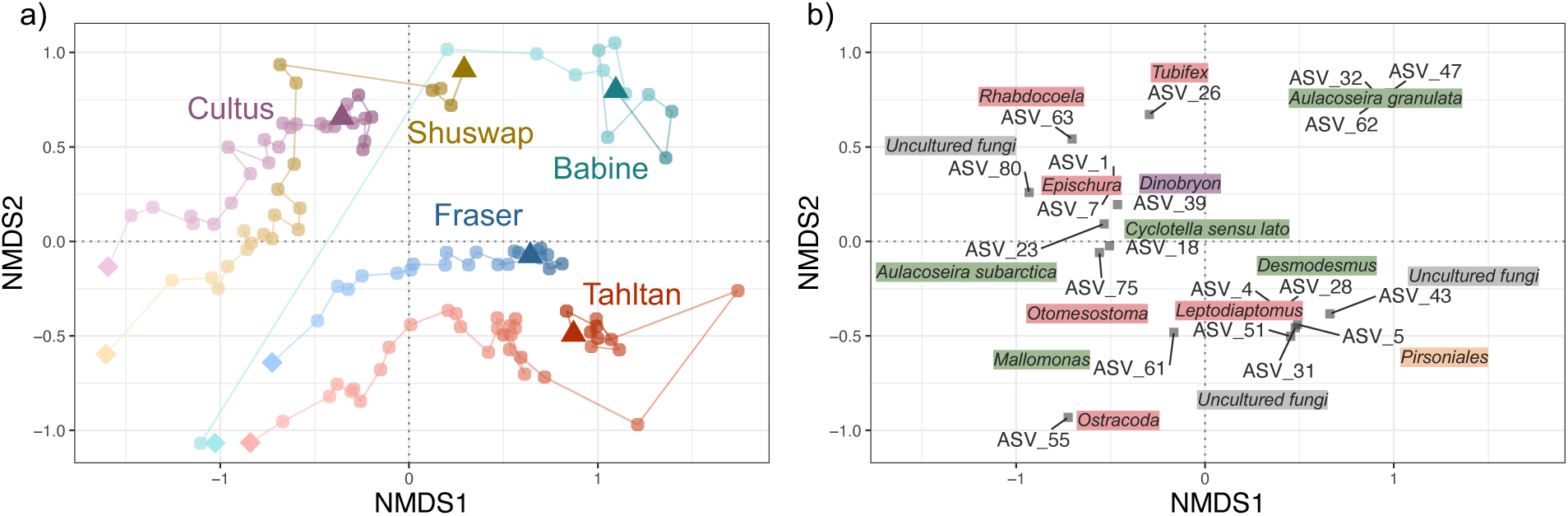
NMDS ordination plots illustrating patterns in salmon nursery lakes. a) Temporal shifts in community composition across salmon nursery lakes, with the top of the cores represented by a small diamond and the bottom of the cores represented by a triangle. b) Amplicons with high abundance and strong loadings passively overlaid. ASVs 32, 47, and 62 were identified as *Aulacoseira granulata*, ASVs 1 and 7 as *Epischura*, and ASVs 4 and 31 as *Leptodiaptomus*.

Lake-specific patterns emerged among invertebrate taxa (Fig. 4b). Cultus and Shuswap lakes exhibited greater sedDNA relative read abundances of *Epischura* (ASV1 and ASV7), while Tahltan Lake was dominated by *Leptodiaptomus* (ASV4 and ASV31). Benthic worms also exhibited lake-specific distributions, with higher relative abundances in Cultus and Shuswap lakes (Fig. 4b). Cultus Lake contained more tubificid worms, particularly *Tubifex* (sludge worm; ASV26) at ∼1976 (8.75cm depth) and *Rhabdocoela* (flatworm; ASV63) at ∼1850s (22.25m depth). Shuswap Lake showed greater abundances of *Otomesostoma* (flatworm; ASV75), especially at ∼1988 (9.5cm depth).

Several amplicons (ASV43, ASV51, ASV80) played important roles in differentiating biological composition both among lakes and through time (Fig. 4b) but had limited taxonomic information. BLASTn assigned them to uncultured fungi with 100% identity. These taxa warrant further investigation to better understand their ecological significance.

### Environmental drivers of community changes with LakePulse dataset integration

To enable broader environmental interpretation, we integrated our data with the LakePulse 18S water column project (Garner et al. 2022; British Columbia subset, *n* = 98; Fig. S10a). Because passive plotting in ordination requires matched variables across datasets, we filtered our sedDNA ASVs to retain only those also present in the LakePulse dataset, preserving on average 63 ± 16% (SD) of the original sedDNA sequence reads. After this filtering, the NMDS trends remained consistent with the original sedDNA trajectories, suggesting the observed patterns with the reduced dataset are robust (Fig. S10b). The most abundant non-overlapping taxa were predominantly green algae (*Desmodesmus*), which were historically more abundant in these salmon nursery lakes, and dinoflagellates (Suessiales), most abundant in the northern remote systems (Babine, Tahltan) which are underrepresented in LakePulse sampling (Table S2).

Environmental variables from the LakePulse dataset were passively plotted alongside the data from our five lakes, to explore potential associations with varying environmental conditions (Fig. 5a). Statistically significant variables included total phosphorus (TP), total nitrogen (TN), and dissolved organic carbon (DOC), which aligned with the top right quadrant (Fig. 5a). The ice-free period (ice_disappear) and lake depth were negatively associated with axis 1 (Fig. 5a). Major ions such as Na^+^, K^+^, and Mg^2+^ were more negatively associated with axis 2, whereas the

**Fig. 5.**
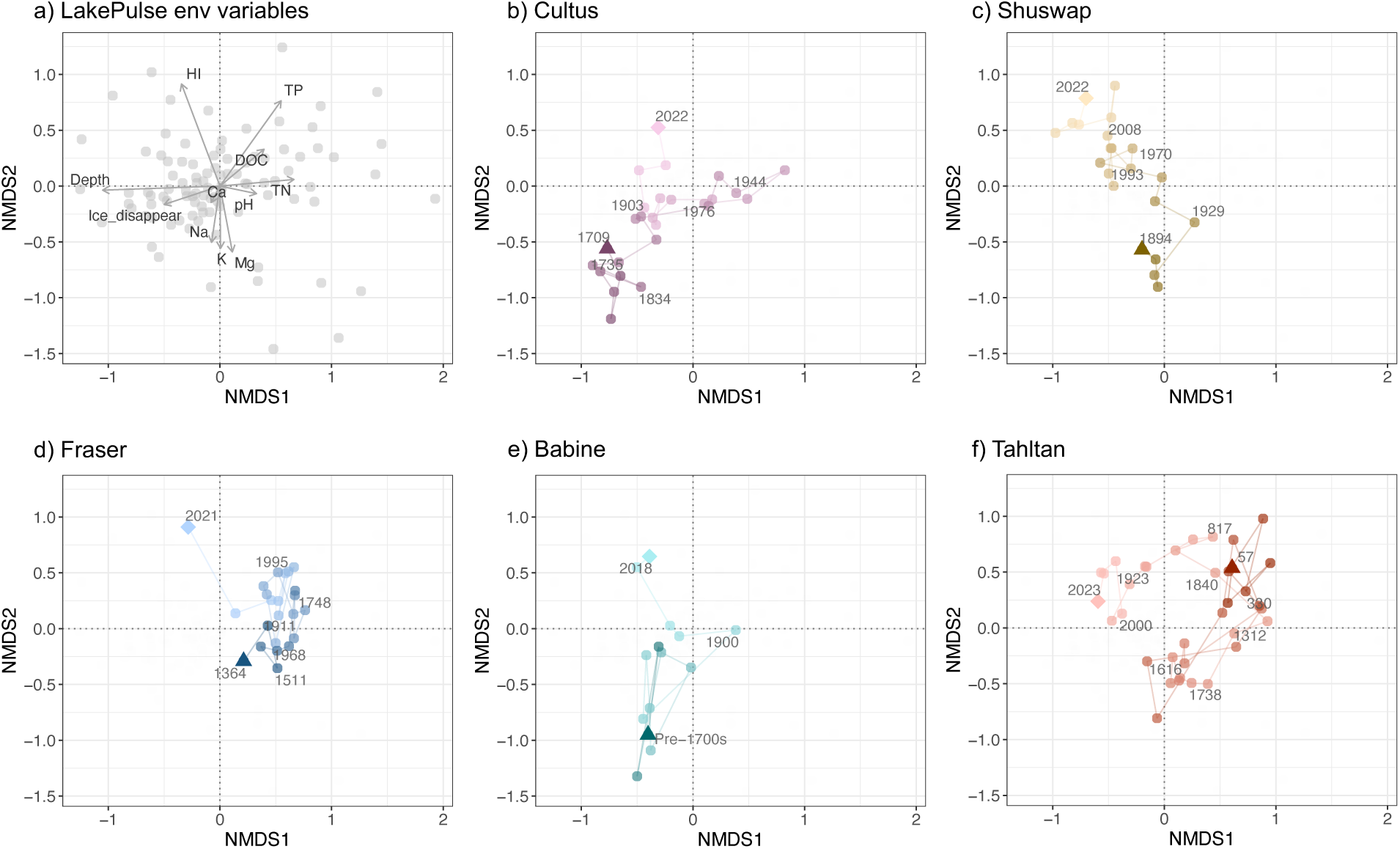
NMDS ordination plots illustrating patterns in the LakePulse BC dataset, with environmental variables and salmon nursery lakes passively plotted. a) Environmental variables: human impact index (HI), total phosphorus (TP), dissolved organic carbon (DOC), total nitrogen (TN), major ions (Ca, Na, K, Mg), ice-off date, and lake water depth. b–f) Passive ordinations of individual salmon nursery lakes: b) Cultus, c) Shuswap, d) Fraser, e) Babine, and f) Tahltan. Dark-colored triangles represent bottom samples; light-colored diamonds represent surface samples.

Human Impact index (HI), reflecting differences in modern land-use among sites, was positively associated with axis 2 (Fig. 5a).

The ordination trajectories revealed distinct temporal patterns in environmental change across the five study lakes (Fig. 5b-f). Cultus Lake (Fig. 5b) initially occupied a position characterized by low nutrient concentrations and low human impact (HI). Around the mid-20th century, its trajectory was more closely associated with higher nutrient and DOC levels, followed by continued movement in recent decades toward regions indicating increased HI. Shuswap Lake (Fig. 5c) displayed a more linear and unidirectional trajectory. Beginning in the 1890s near the lower end of the second axis, samples gradually shifted upward toward positions related to higher HI over time. Fraser Lake (Fig. 5d) showed minor fluctuations along the first axis throughout much of its record, with the most recent samples indicating an abrupt transition toward positions related to higher HI values. Babine Lake (Fig. 5e) began in the bottom left quadrant, associated with low nutrients. Around the 1900s, its trajectory gradually shifted toward higher nutrient and DOC conditions, followed by recent movement toward positions related to elevated HI. Tahltan Lake (Fig. 5f) exhibited the most dynamic trajectory. It initially occupied a position characterized by high nutrient levels, then moved toward greater ion concentrations during a period marked by a distinct change in sediment lithology (Details see Supporting Information S2). Around the 1840s, it returned to a more nutrient-rich state, followed by a leftward shift, associated with a longer ice-free season over the past century.

### Co-occurrence network analyses

Network analyses of co-occurrence patterns among dominant ASVs, applied to zones of community compositional change identified through CONISS, revealed substantial restructuring across all five lakes (Fig. 6 for Cultus and Tahltan; Fig. S11 for Shuswap, Fraser, and Babine). The functional composition of these dominant ASVs, categorized as heterotrophs, mixotrophs, parasites, and phototrophs, shifted across temporal zones (Fig. 6 & Fig. S11). In Cultus Lake, heterotroph relative read abundances increased from Zone 1 to Zone 2 (defined based on a constrained cluster analysis), then phototrophs increased from Zone 2 to Zone 3 (Fig. 6a).

**Fig. 6.**
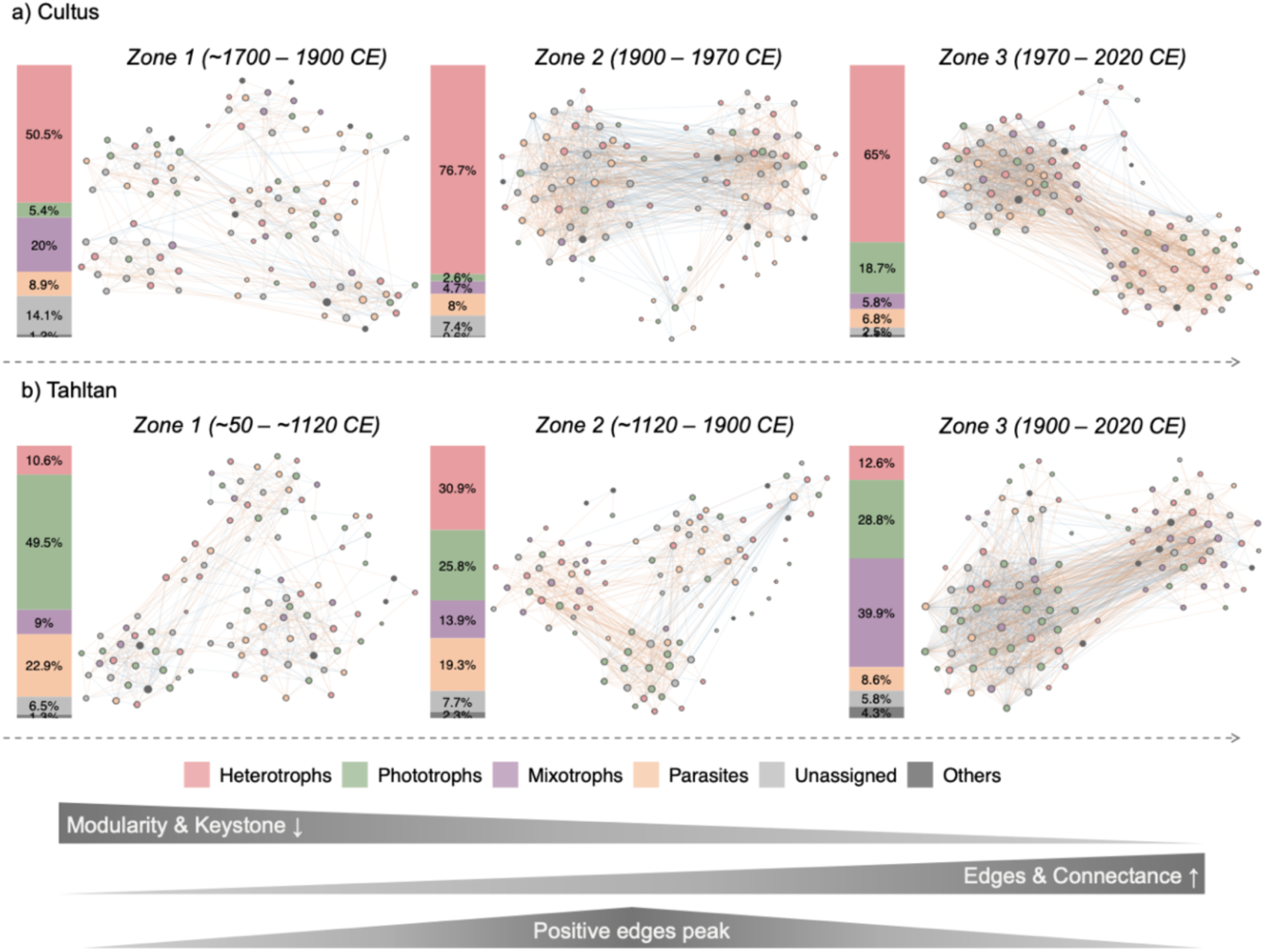
Network analysis illustrating co-occurrence patterns of dominant ASVs in a) Cultus and b) Tahltan lakes across distinct time zones. Red edges indicate positive correlations while blue edges represent negative associations. Overall trends in network modularity, total edges, node connections, and the proportion of positive edges from the two focal lakes are displayed in the lower panels. Bar plots show the relative proportions of dominant amplicons within each zone, classified into five trophic groups.

Tahltan Lake showed a similar increase in relative read abundances of heterotrophs from Zone 1 to Zone 2, but mixotrophs increased from Zone 2 to Zone 3 (Fig. 6b). Shuswap Lake showed increased relative read abundances in phototrophs from Zone 1 to Zone 2 (Fig. S11a), whereas Fraser Lake showed increased relative read abundances in parasites and mixotrophs (Fig. S11b), and Babine Lake showed increased heterotrophs and phototrophs (Fig. S11a).

The co-occurrence network structure also changed over time, with modularity decreasing while total edges and node connections increased (Figs. 6, & Fig. S11). The proportion of positive edges peaked during Zone 2 in both Cultus and Tahltan lakes. Historical networks exhibited a highly modular structure across all lakes, with communities organized into multiple discrete modules characterized by distinct, well-separated clusters of taxa and relatively few connections between them (Fig. 6 & Fig. S11). As time progressed, networks became increasingly interconnected and less modular across all sites, accompanied by a general increase in the number of edges (Fig. 6 & Fig. S11).

The relative distribution of the topological roles in the network (i.e., connectors, module hubs, network hubs, and peripheral nodes) also changed across the zones (Fig. S12 & S13).

Historical networks contained more module hubs and network hubs, reflecting their highly modular organization with well-defined keystone taxa. In contrast, modern networks were dominated by peripheral nodes with only a few remaining hubs, consistent with the shift toward less modular, more interconnected community structures.

## DISCUSSION

Historical changes in eukaryotic communities across our study lakes provide insights into past ecosystem dynamics driven by both natural and anthropogenic forces. The sedDNA time series reveal significant shifts in algal community composition over the past few centuries to millennia, and reflect how these salmon nursery lake habitats may have responded to changes in water column mixing regimes and nutrient dynamics. Beyond taxonomic shifts, we explored ecosystem structure and potential species interactions using co-occurrence network analyses, offering a community-level perspective on ecological change. We synthesized the limnological histories of Cultus and Tahltan Lakes by integrating temporal patterns, ordination trajectories, and network structure (Supplementary Information S2). Finally, we acknowledged key limitations of sedDNA metabarcoding and outlined future research directions to improve ecological inference from paleogenetic records.

### Long-term algal dynamics in Pacific salmon nursery lakes

The sedDNA assemblages of each algal group exhibited distinct temporal shifts. We interpreted these shifts of major phytoplankton taxonomic groups in the context of different environmental drivers.

### Bacillariophyta

A notable trend across four of the five study lakes was the increasing relative abundances of diatom reads, marking a major shift in algal community structure (Fig. 2a). These increases often followed a two-phase pattern: an initial rise in benthic (e.g., *Staurosira*) and tychoplanktonic (e.g., *Aulacoseira*) taxa, followed by the recent dominance of planktonic forms within the *Cyclotella* sensu lato (Fig. 2b). This trajectory broadly aligns with patterns previously documented in subfossil diatom records (Rühland et al. 2015; Michelutti et al. 2020).

The initial phase likely reflected a longer ice-free season, during which nearshore zones received more sunlight and heat, promoting benthic diatom growth (Smol 1988). As mixing increased across the water column, tychoplanktonic taxa such as *Aulacoseira* benefited. The recent surge in planktonic diatoms was potentially driven by earlier spring ice-off, which could have extended the spring isothermal mixing period before summer stratification. This provide competitive advantages to small planktonic taxa that thrive under well-mixed spring conditions, as supported by monitoring data from lakes in the Experimental Lakes Area (ELA, Ontario, Canada), showing that *Discostella* (a planktonic centric diatom genus) is often more abundant during spring and early summer (April-June; Wiltse et al. 2022). Similar spring maxima for *Discostella* have been documented in other BC lakes where winter has had greater temperature increases compared to other seasons (Laird et al. 2021).

Among our study lakes, three years of monthly monitoring data from Cultus Lake (Gauthier et al. 2021) using the same primer set provided additional evidence for enhanced relative abundances of centric planktonic diatoms (i.e., *Cyclotella* sensu lato) occurring during an extended mixing period (Fig. 7). From July 2014 to June 2017, these planktonic diatom reads consistently peaked during winter and spring months rather than summer. As an ice-free monomictic system, Cultus Lake undergoes mixing from late fall (November-December) to early May. During these prolonged winter and spring mixing periods, total phosphorus (TP) concentrations remained more elevated (epilimnetic TP: 8.4 μg/L during mixing vs. 5.4 μg/L during stratification) and uniformly distributed throughout the water column (Gauthier et al. 2021), creating conditions that enhanced diatom productivity and favored the growth of these planktonic taxa.

**Fig. 7.**
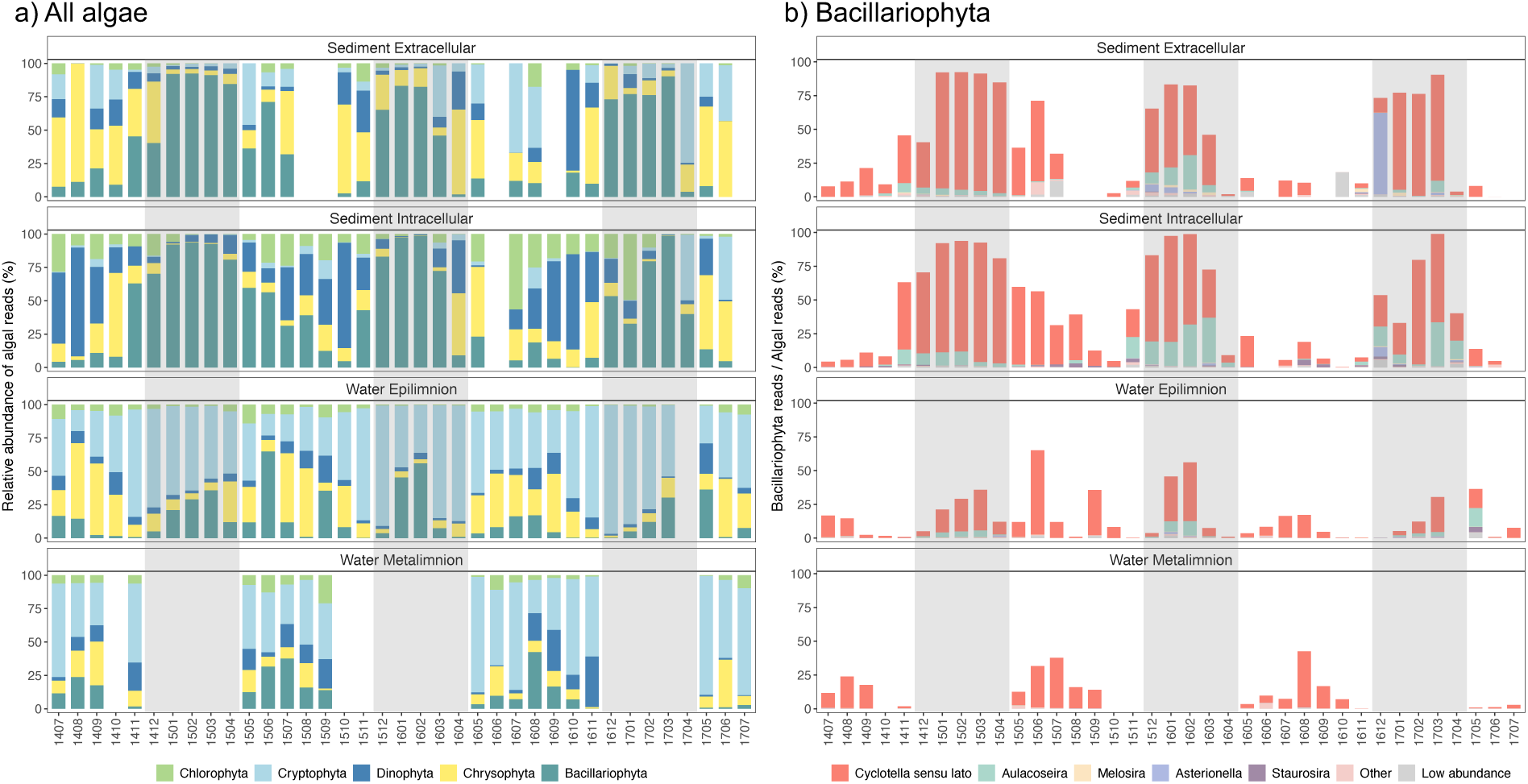
Algal community composition and diatom abundance in Cultus Lake based on 36 months of sampling. (a) Temporal dynamics of major algal groups. (b) Relative abundance of diatoms as a proportion of total sequence reads. Grey shaded areas indicate periods of water column mixing.

Babine Lake exhibited distinct diatom assemblage dynamics compared to the other study lakes, which may be explained by its unique morphometry. For much of the sedDNA record, Babine Lake was dominated by tychoplanktonic *Aulacoseira*. This persistent dominance can be linked to the elongated shape of Babine Lake and long wind fetch - features that promote strong and sustained water column mixing. Such conditions are ideal for *Aulacoseira*, which thrives in turbulent environments. However, post-1950s, a regional decline in wind speeds may have weakened mixing (Barouillet et al. 2024), contributing to a decline in *Aulacoseira* abundance. At the same time, longer ice-free seasons likely promoted the growth of benthic taxa like *Staurosira*, whose relative abundance has increased since the mid-20th century (Fig. 2b).

### Chlorophyta

The systematic decline of green algae across all study systems represented one of the most consistent patterns in our dataset (Fig. 2). The magnitude of this decline was particularly striking in Fraser and Tahltan lakes, where green algae relative abundance decreased from >70-90% historically to <3% in recent samples. This decline was likely due to enhanced thermal stratification in recent decades, which created prolonged nutrient depletion in surface waters.

Chlorophytes generally thrive in well-mixed, nutrient-rich environments (Celewicz et al. 2022). As primarily non-motile, obligate photoautotrophs, they are largely restricted to the epilimnion, where light is sufficient, but nutrient availability declines progressively during periods of stratification.

### Chrysophyta

Chrysophytes showed increasing trends across most study lakes over the past century, consistent with other paleolimnological records (Wolfe and Silver 2013; Mushet et al. 2017). Two main factors may explain these increases: (1) rising lakewater pCO₂ and (2) enhanced thermal stratification from regional warming.

Unlike many phytoplankton, chrysophytes lack carbon-concentrating mechanisms (CCM) for converting HCO₃⁻ to CO₂, because they evolved when aquatic CO₂ was more abundant (Raven et al. 2012). Therefore, they rely heavily on dissolved CO₂ for photosynthesis (Wolfe and Silver 2013), explaining their prevalence in low-pH lakes where CO₂ remains gaseous rather than converted to bicarbonate. Rising lakewater pCO₂, possibly due to increasing atmospheric CO₂ concentrations (i.e., higher equilibrium CO₂ levels in surface waters) and/or enhanced mineralization of organic matter within lakes (Wolfe and Silver 2013), may therefore favor chrysophyte dominance.

With enhanced regional warming (Fig. 1b), chrysophytes likely benefit from enhanced thermal stratification through their motility, which allows them to optimize their position within the water column to access favorable temperature, nutrient, and light conditions (Heinze et al. 2013). Additionally, some chrysophytes (e.g., *Ochromonas* & *Dinobryon*) are mixotrophic, enabling them to supplement photosynthesis with phagotrophy when light is limited in deeper waters (Lie et al. 2017). This combination of motility and nutritional flexibility likely resulted in the chrysophyte blooms in the hypolimnion during stratified periods (Fee 1976). The 36-month monitoring record from Cultus Lake revealed elevated chrysophyte relative abundances during stratified periods, particularly in summer (Fig. 7a), when mixotrophic chrysophytes can be highly competitive.

### Dinophyta

Dinoflagellates increased in Shuswap Lake after the 1950s and in Fraser and Tahltan Lakes after the 1900s (Fig. 4b), possibly reflecting enhanced thermal stratification as well. As primarily mixotrophic organisms, dinoflagellates supplement photosynthesis through bacterial ingestion and use vertical migration to access light in surface waters and nutrients at depth (Stoecker et al. 2017; Zheng et al. 2023). Based on the seasonal monitoring data of Cultus Lake, we confirmed that dinoflagellates reached their peak during stratified periods, when they often exceeded 50% of algal sedDNA sequences (Fig. 7a). The more recent declines in the relative read abundance of dinoflagellates in Cultus, Shuswap, Fraser and Tahltan lakes may reflect disproportionate increases in other algal groups such as diatoms and chrysophytes, but quantitative PCR analyses would be needed to confirm this hypothesis.

### Cryptophyta

Cryptophytes were consistently detected at low levels across all study sites. This may reflect (1) genuinely low abundance, or (2) could be indicative of high grazing pressure due to their palatability (Tonno et al. 2016), contributing to lower cellular deposition compared to other taxa, or (3) due to poor preservation due to their lack of rigid cell walls (Hoef-Emden and Archibald 2016). These latter hypotheses are supported by Cultus Lake 36-month monitoring, where cryptophytes dominated water column samples (both epilimnion and metalimnion; Fig. 7a) but were underrepresented in sediment traps, suggesting intense grazing in the water column and/or higher degradation while sedimenting (Gauthier et al. 2021). Additionally, higher cryptophyte proportions in the extracellular versus intracellular sedDNA fraction (Fig. 7a) likely indicate their lack of protective walls renders them vulnerable to degradation in sedDNA pool.

### Temporal trends in other taxa

Copepods dominated zooplankton assemblages across our study lakes (Fig. S8), consistent with their documented prevalence in BC Pacific salmon nursery lakes, where they may often comprise more than half of the zooplankton communities (Self and Larratt 2021). *Epischura* dominated the zooplankton sedDNA reads in Cultus, Shuswap, and Fraser Lakes, while *Leptodiaptomus* dominated in Tahltan Lake (Fig. 4b & Fig. S8). Both genera are relatively large-bodied calanoid copepods (Perez-Fuentetaja et al. 1996): *Epischura* consume algae in early life stages and cladocerans and other copepods as adults (Chow-Fraser and Wong 1986), while *Leptodiaptomus* species are primarily herbivorous (Torke 2001). Unfortunately, species-level identification cannot be achieved due to high sequence similarity at this marker across copepods; otherwise, more detailed species-specific autecology could be used to infer their precise roles in the food web. Rotifers and cladocerans appeared infrequently and at low abundances in the sediment records. This pattern likely reflects either their naturally lower biomass relative to copepods in the water column, reduced preservation of their remains in sediments (e.g., rotifers may degrade more rapidly than copepods), and/or limited amplification efficiency for these taxa using the 18S rRNA V7 marker. For instance, *Daphnia* were absent from our metabarcoding results despite being recorded in high abundance via subfossil remains in our lakes (Gauthier et al. 2020; Barouillet et al. 2024), likely due to their exceptionally long 18S rRNA gene V7 regions (Crease and Colbourne 1998).

The successful detection of multiple fish species (Fig. S9), including *Oncorhynchus nerka* (sockeye salmon), demonstrates the potential of sedDNA to reconstruct fish community dynamics, despite challenges associated with degraded DNA, particularly for high-trophic-level taxa with low overall biomass (Huston et al. 2023). The predominance of sockeye salmon sequences relative to other fish species is consistent with the established role of these lakes as salmon nursery habitats. The generally declining trends in fish DNA across most lakes contrast with the increasing trend observed in Babine Lake (Fig. S9), aligning with expectations from the Babine Lake Development Project (BLDP; Cox-Rogers and Spilsted 2012). This divergence suggests that while many systems may be experiencing fish population fluctuations or declines due to anthropogenic stressors, Babine Lake may be showing a positive response to management interventions. However, fish DNA read counts were generally low across all lakes, and these trends represent proportional changes relative to other eukaryotic organisms rather than absolute abundances. If algal and zooplankton biomass increased more substantially than fish biomass, fish DNA would exhibit a declining trend even if absolute fish DNA quantities remained stable or increased slightly. Nevertheless, the detection of fish DNA in sediment cores represents a promising avenue for exploring alternative primers and molecular approaches to detect and quantify fish DNA in paleolimnological studies for management purposes.

### Co-occurrence of eukaryotic taxa

Network analysis revealed a fundamental transformation in community organization across all study lakes, shifting from highly modular structures to increasingly connected but less modular networks over time (Fig. 6 & Fig. S11). The pronounced modularity in the earliest zones likely reflects stable communities with distinct taxa clusters and minimal inter-module interactions, a configuration that buffers perturbation spread by containing disturbances within compartments (Stouffer and Bascompte 2011; Gilarranz et al. 2017).

Over time, increasing connectance and declining modularity may indicate a shift toward more generalized biotic interactions and greater functional redundancy. This was evident in taxonomic expansion within functional groups: for example, chrysophyte assemblages shifted from single-genus dominance in historical samples to multi-genus representation in recent samples (Fig. 3a). Multiple species may now occupy similar ecological roles within each nutritional guild, creating insurance against species loss through functional redundancy - where the loss of one species can be compensated by ecologically similar taxa (Biggs et al. 2020).

The peaks in positive connections during the middle time zones in both Cultus and Tahltan lakes suggest important transitions (Fig. 6), despite different environmental histories. Increases in positive associations could reflect environmental disturbances eroding competitive hierarchies and niche specialization, allowing previously excluded species to coexist via facilitation or reduced niche overlap (Hernandez et al. 2021; McMeans et al. 2020). For example, warming can alter thermal stratification, compressing phytoplankton into narrower depth ranges and increasing spatial overlap that promotes coexistence by preventing single-species dominance (Wang et al. 2024). The transitional phases marked the breakdown of modular compartments in Zone 1 and the establishment of new coexistence patterns that facilitate the buildup of functional redundancy observed in Zone 3 (Fig 6).

The prevalence of mixotrophs in recent intervals (Tahltan and Fraser lakes; Fig. 6b & Fig. S11b) was particularly noteworthy, as mixotrophy confers resilience through metabolic versatility, enabling organisms to switch between photosynthetic and heterotrophic strategies under fluctuating conditions (Stoecker et al. 2017; Floder et al. 2024). This network reorganization may represent a fundamental shift in resilience mechanisms: from compartmentalized specialists buffering between modules to functional overlap among generalists providing insurance within guilds.

The progressive loss of module hubs and network hubs from historical to modern zones (Figs. S12, S13) suggests declining numbers of keystone taxa that previously structured these communities. Similar declines in hub taxa and modularity under environmental stress have been documented in other ecosystems (Hernandez et al. 2021; Wang et al. 2025), highlighting that this may be a common response to anthropogenic pressures. This hub-to-peripheral transition reflects a fundamental reorganization of community structure that may compromise resilience to future perturbations.

### Limitations and future perspectives

The 18S rRNA V7 gene marker has allowed us to reconstruct lake community dynamics over centennial to millennial timescales. Our analysis revealed diverse algal assemblages, calanoid copepods, and a range of parasites associated with both algae and invertebrates. While fish DNA was detected, its occurrence was sporadic and at low abundance (Fig. S9), which may reflect both the relatively low fish biomass in the lake and methodological limitations, as DNA degradation and PCR amplification biases can disproportionately affect the recovery of vertebrate DNA from complex sedDNA pools. This suggests that more targeted approaches, such as deep-sequencing metagenomics may be necessary to adequately characterize these high-trophic, low-biomass organisms.

The taxonomic resolution of the 18S rRNA gene V7 region presents limitations for certain groups. For example, we were unable to resolve dinoflagellates and rotifers to the genus level due to both high sequence similarity among taxa and incomplete representation in reference databases, which are particularly lacking for regionally distinct or less-studied organisms.

Researchers specifically interested in these taxa may benefit from applying more targeted, group-specific markers to achieve higher taxonomic resolution. Nonetheless, for our broader ecological objectives, 18S rRNA V7 marker provided a reasonable balance between coverage and resolution across a wide diversity of eukaryotic taxa.

Cladocerans remain particularly challenging in sedaDNA studies. The 18S rRNA gene V7 region of *Daphnia* is unusually long, potentially leading to poor recovery using the 2 × 250 bp Illumina sequencing platform. We tested two alternative markers - Cytochrome Oxidase subunit I (COI; mlCOIintF-jgHCO2198) and the 18S rRNA gene V9 region (1391F - EukBr; data not shown) - but neither proved excellent for cladoceran detection in sediment cores. COI fragments (∼313 bp) are typically too long for degraded sedDNA, and while the 18S rRNA gene V9 region yielded data, much of it comprised bacterial sequences or was dominated by algae and copepods (data not shown). Future research targeting cladocerans in sedDNA archives will likely require the development and validation of dedicated group-specific primers and protocols.

## CONCLUSIONS

This study examined long-term ecological changes in five salmon nursery lakes in British Columbia using sedimentary DNA metabarcoding, revealing major shifts in primary producers and consumers over the past centuries to millennia. Algal communities showed consistent increases in the relative abundance of planktonic diatoms (*Cyclotella* sensu lato), and seasonal monitoring data revealed these taxa dominate during winter-spring mixing. Green algae declined sharply, likely due to competitive disadvantage under stratified conditions where surface waters become nutrient-depleted, while chrysophytes and dinoflagellates generally increased in abundance, benefiting from their motile and mixotrophic capabilities under enhanced thermal stratification over the past century.

Such community shifts have implications for salmon nursery function. The increase in diatoms like *Cyclotella* sensu lato may enhance zooplankton food quality because they typically supply essential fatty acids (e.g., EPA) to grazers (Brett & Müller-Navarra 1997), but the concurrent rise of grazing-resistant mixotrophic flagellates (e.g., scaled chrysophytes) may counteract these benefits by reducing grazing efficiency (Vad et al. 2020). Cladocerans, particularly *Daphnia*, are the primary prey of juvenile sockeye salmon (Hampton et al. 2006), yet our 18S rRNA gene V7 primer set failed to capture them, preventing assessment of whether changes in primary producers are cascading to this critical trophic link. Although fish DNA was detected across all lakes, read counts were low, limiting our ability to infer population trends from these data alone. Nonetheless, the inferences drawn from the sedDNA assemblages about changes in thermal mixing patterns of the nursery lakes could have important implications for the development of juvenile salmon, as regional temperature has been previously identified as an important predictor of early growth patterns, though its effects are modulated by local characteristics (Price et al. 2024).

The co-occurrence network analysis revealed a shift from highly modular communities with distinct functional compartments to more connected assemblages with greater functional redundancy, representing a fundamental change in resilience mechanisms from compartmentalized specialists to functionally redundant generalists. Lake-specific history reveals that Cultus Lake was shaped by human activity beginning in the late 19th century, including logging, recreation, and shoreline development, while Tahltan Lake displayed complex responses likely due to a combination of climate and/or hydrological changes and shifts in nutrient loading from salmon and terrestrial inputs. In both lakes, declining network modularity and rising positive interactions signaled major ecosystem reorganization.

Despite differences in geographical and historical background, all study lakes converged toward higher human impact and/or longer growing seasons in recent decades. This shared trajectory underscores the widespread and accelerating influence of human-induced watershed disturbance and climate change on aquatic ecosystems. Our findings demonstrated that sedDNA can effectively reconstruct ecological histories, though its limitations (e.g., taxonomic resolution and preservation biases) require cautious interpretation. Overall, these results highlight the urgent need for more adaptive and integrated freshwater management strategies in the face of mounting environmental change. For example, the Babine Lake sedDNA profile appears to have captured both variation in climate dynamics and potential responses to fisheries management. As sedDNA analysis remains a developing method, integration with complementary paleoecological indicators and additional molecular or traditional approaches is recommended.

