## Supplementary Information for "Changing Pacific salmon nursery lake ecosystem dynamics over centuries to millennia: insights from sedimentary DNA metabarcoding"

### SUPPORTING INFORMATION

#### Supporting Information S1. Local History

Local history varies considerably across our study sites, with Fig. S1 outlining major events related to human settlement, land-use development, and environmental changes that reveal distinct temporal patterns of anthropogenic influence across each lake. **Cultus Lake** shows the most intensive modern development, beginning with Euro-American settlement in 1880, followed by road construction (1900), logging peaks (1920), and establishment as a provincial park (1948). Recent decades have seen extensive recreational development (1990s), invasive species introductions (Eurasian watermilfoil, 1977), and ongoing fisheries management challenges including sockeye population endangerment (2003) and fishery closures (2004). **Shuswap Lake** exhibits a similar southern development pattern, with European settlement beginning in 1860, railway access (1885), and town establishment (1905). The timeline shows continued agricultural and recreational development through the 20th century, with recent environmental pressures (forest fire) and ongoing sockeye population fluctuations. **Fraser Lake** demonstrates more modest development pressure, with key events including fur-trading post establishment (1800), village incorporation (1966), and molybdenum mining (1966). Recent decades show mining suspension (2015), alongside environmental changes including cyanobacterial blooms (2016) and fire events (2018, 2024). **Babine Lake** reflects industrial resource extraction impacts, particularly large-scale deforestation (1900-1920) and overfishing (1910), followed by major infrastructure development (Babine Lake Development Project, 1962). Recent management efforts focus on habitat restoration, including mine closures (1990) and enhanced salmon spawning programs. **Tahltan Lake** has been home to the Tahltan Nation for millennia, with initial European contact occurring through Indigenous trade networks (Dease

Lake trading post, 1837) followed by increased numbers of people passing through the village during the Klondike Gold Rush (1898). Modern impacts remain minimal, with recent changes primarily related to improved access (highway completion, 1972) and temporary mining operations (1952-1992).

Supporting Information Table S1. Weather station information: including ID, distance to our study lakes, records starting and ending year.

| Lake | Station | Station_ID | Distance_km | Year_start | Year_end |
|---|---|---|---|---|---|
| Cultus | Chilliwack | 735 | 15 | 1879 | 2014 |
| Shuswap | Westwold | 1344 | 99 | 1921 | 2013 |
| Fraser | Fort St James | 588 | 66 | 1895 | 2019 |
| Babine | Smithers A | 487 | 131 | 1942 | 2018 |
| Tahltan | Dease Lake | 1454 | 186 | 1944 | 2011 |

Supporting Information Table S2. Top 10 most abundant ASVs in salmon lake sedDNA records without overlap in the LakePulse dataset. ASVs are ranked by total read counts across all sediment samples. Taxonomic classifications shown from Division to Genus level; blank cells indicate unresolved taxonomy.

| **Amplicon** | **Total reads** | **Prop (%)** | **Division** | **Class** | **Order** | **Genus** |
|---|---|---|---|---|---|---|
| ASV_8 | 177574 | 1.91 | Chlorophyta | Chlorophyceae | Sphaeropleales | Desmodesmus |
| ASV_13 | 107274 | 1.15 | Alveolata | Dinophyceae | Suessiales | |
| ASV_14 | 105571 | 1.14 | Chlorophyta | Chlorophyceae | Sphaeropleales | Desmodesmus |
| ASV_22 | 66427 | 0.72 | Rhizaria | | | |
| ASV_26 | 57435 | 0.62 | Opisthokonta | Annelida | | Tubifex |
| ASV_27 | 56899 | 0.61 | Rhizaria | | Plasmodiophorida | |
| ASV_28 | 56189 | 0.60 | Chlorophyta | Chlorophyceae | Sphaeropleales | Desmodesmus |
| ASV_29 | 54239 | 0.58 | Stramenopiles | Bacillariophyceae | Fragilariales | Staurosira |
| ASV_30 | 53702 | 0.58 | Opisthokonta | Nematoda | Chromadorea | Eumonhystera |
| ASV_31 | 51806 | 0.56 | Opisthokonta | Arthropoda | Crustacea | Leptodiaptomus |
| ASV_32 | 51122 | 0.55 | Stramenopiles | Coscinodiscophyceae | | |
| ASV_34 | 48607 | 0.52 | Rhizaria | | Plasmodiophorida | |

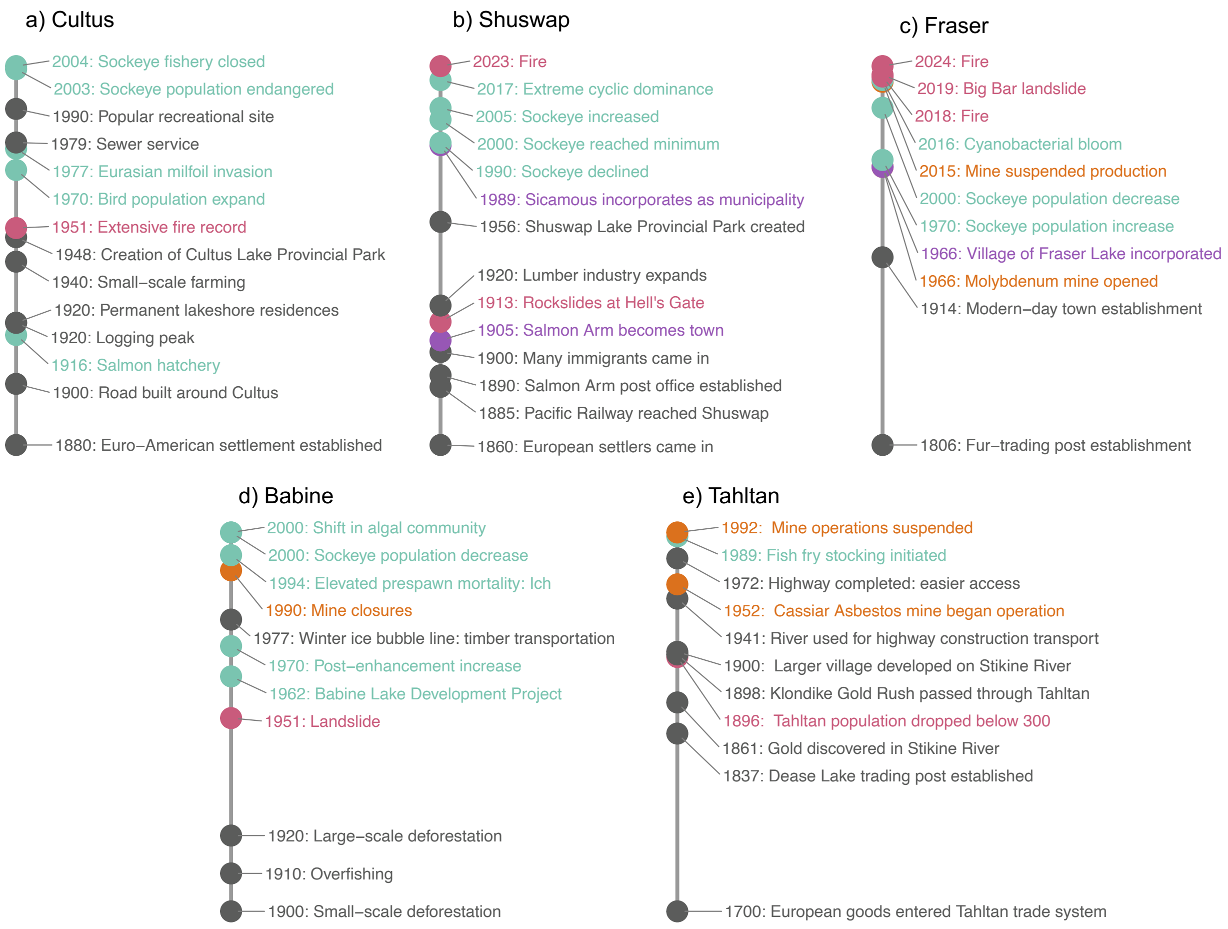


Supporting Information Fig. S1 Historical context of the five salmon nursery lakes, highlighting key events related to development activities (gray), industrial operations (orange), government policies/regulations (purple), natural or anthropogenic disasters (pink), and environmental changes (green) for: a) Cultus, b) Shuswap, c) Fraser Lake, d) Babine Lake, and e) Tahltan Lake.

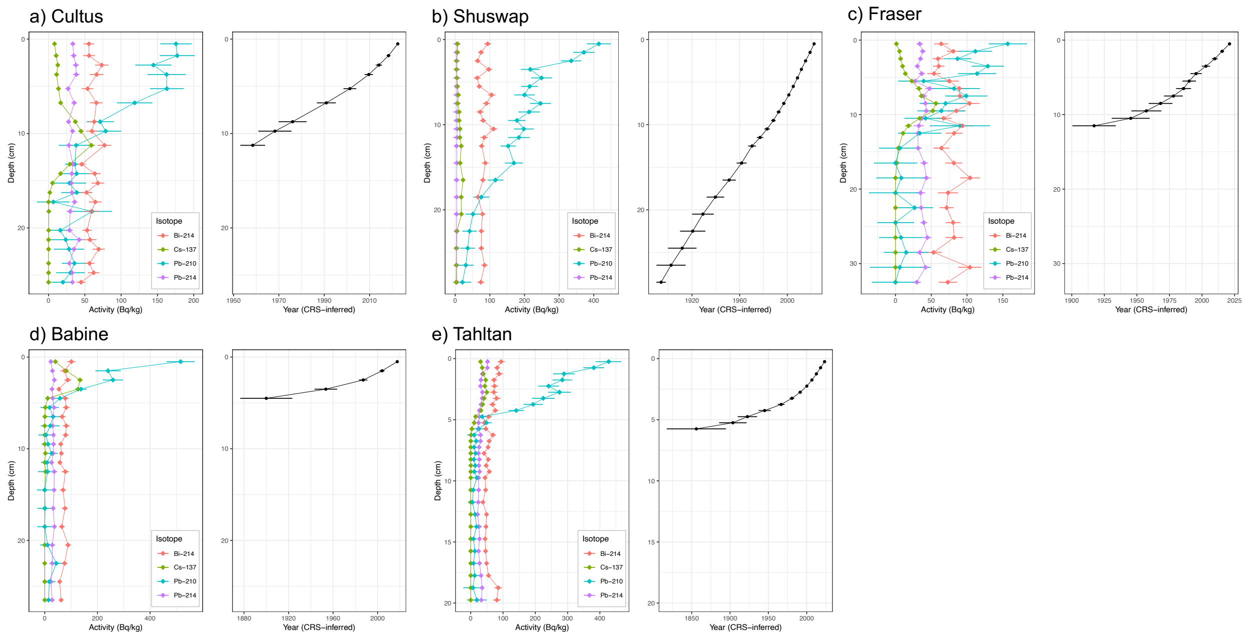


Supporting Information Fig. S2 Isotope activities (Bi-214, Cs-137, Pb-210, and Pb-214) and corresponding constant-rate-of-supply (CRS) age-depth models for: a) Cultus, b) Shuswap, c) Fraser, d) Babine, and f) Tahltan Lake. Error bars indicate uncertainties associated with the CRS model.

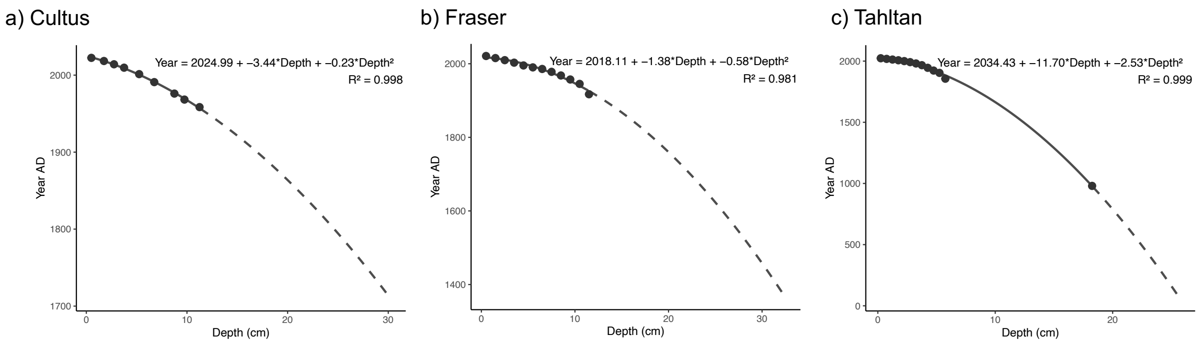


Supporting Information Fig. S3 Polynomial models for (a) Cultus, (b) Fraser, and (c) Tahltan lakes. Radiocarbon dating was performed on two intervals from the core of Tahltan Lake. We use the $^{14}C$ date from the interval at 18.25 cm depth (calibrated to 939-1022 AD at 65.4% probability and 892-933 AD at 30% probability), which yielded an age of approximately 980 AD and corresponds closely with the depth of the C:N peak (~1000 AD) identified by Selbie (2008). A second $^{14}C$ sample at 24.25 cm depth (calibrated to 1116-1219 AD at 61.1% probability and 1042-1087 AD at 29.2% probability) produced an age reversal and was excluded from the age model, as it likely does not represent the true sediment age. All 14C dates were based accelerated mass spectrometry analyses conducted on extracted pollen material by Beta Analytical.

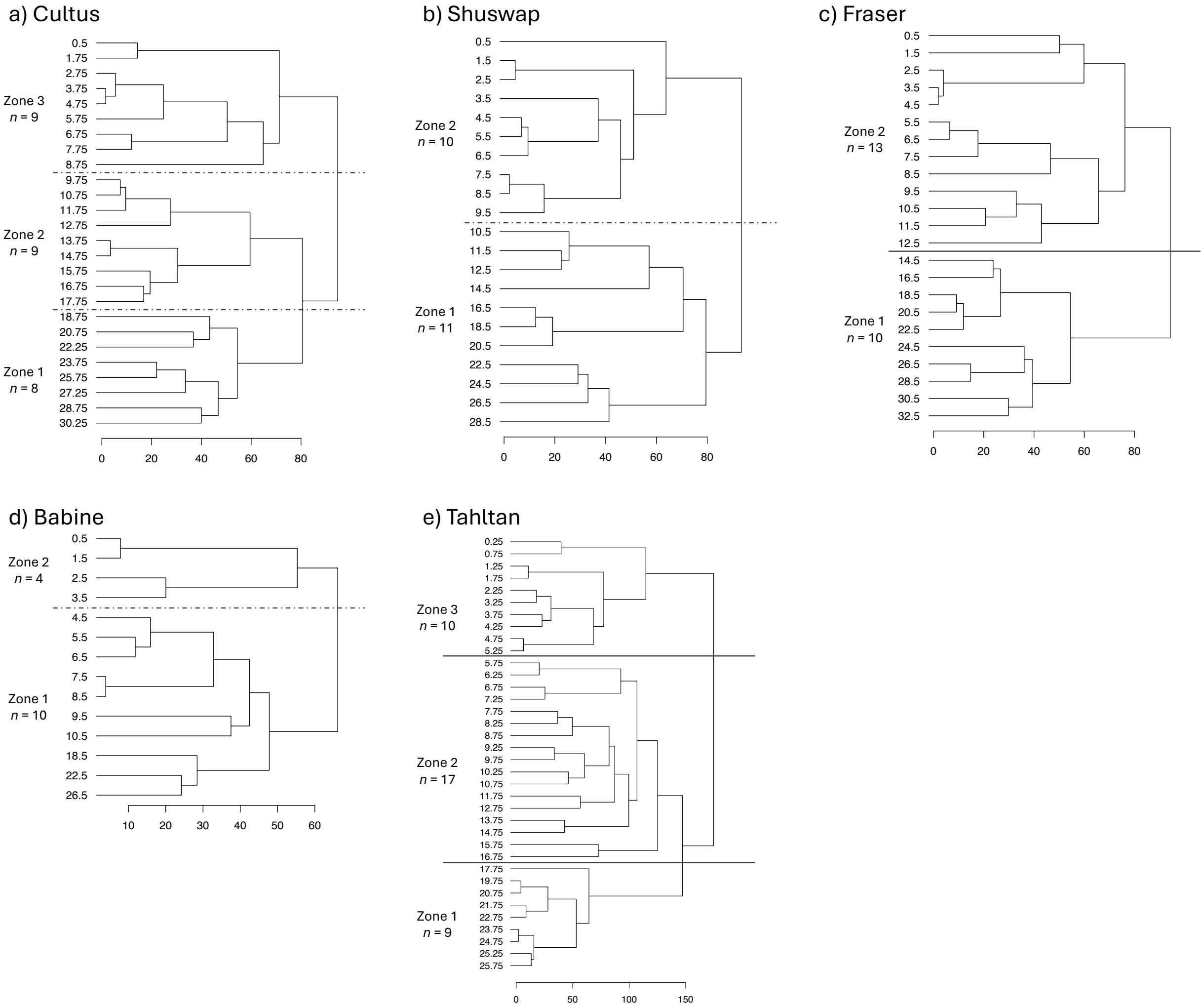


Supporting Information Fig. S4 Stratigraphically constrained cluster analysis (CONISS) for: a) Cultus, b) Shuswap, c) Fraser, d) Babine, and f) Tahltan lakes. Zone divisions are shown in dashed lines (where broken stick model did not yield a distinct zone, we selected the first cluster(s) identified) and solid lines (where broken-stick model shows significant division).

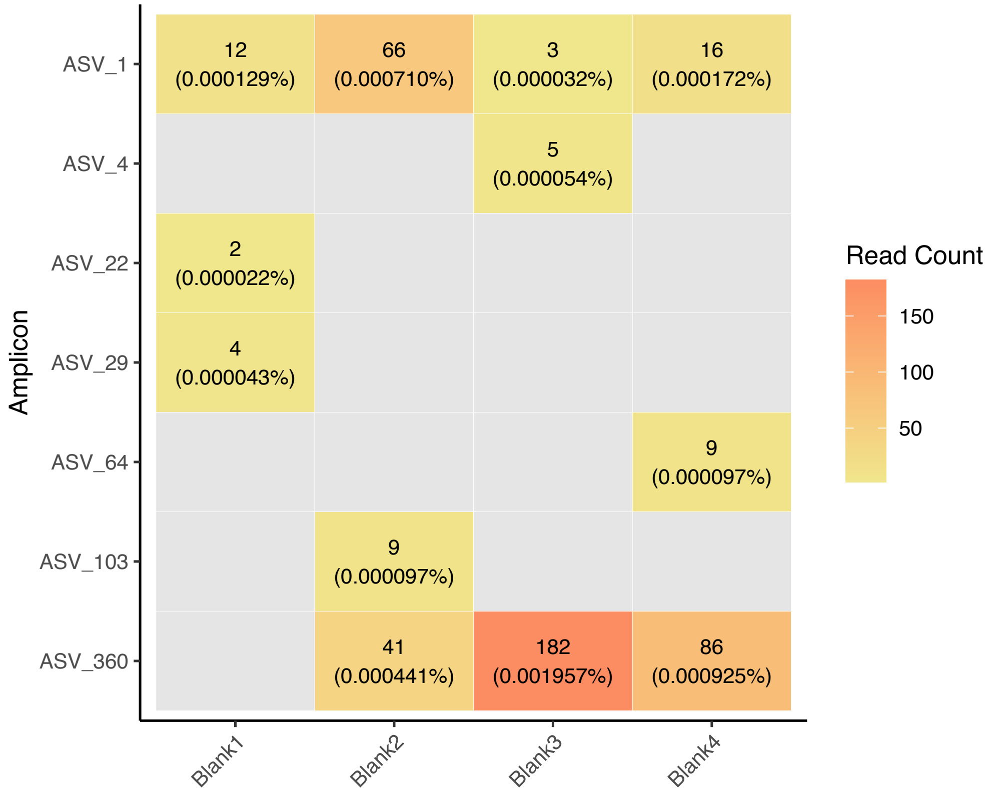


Supporting Information Fig. S5 Heatmap showing the sequence counts and percentages out of total sequences (n = 9,298,026) of each ASV detected in the blank controls.

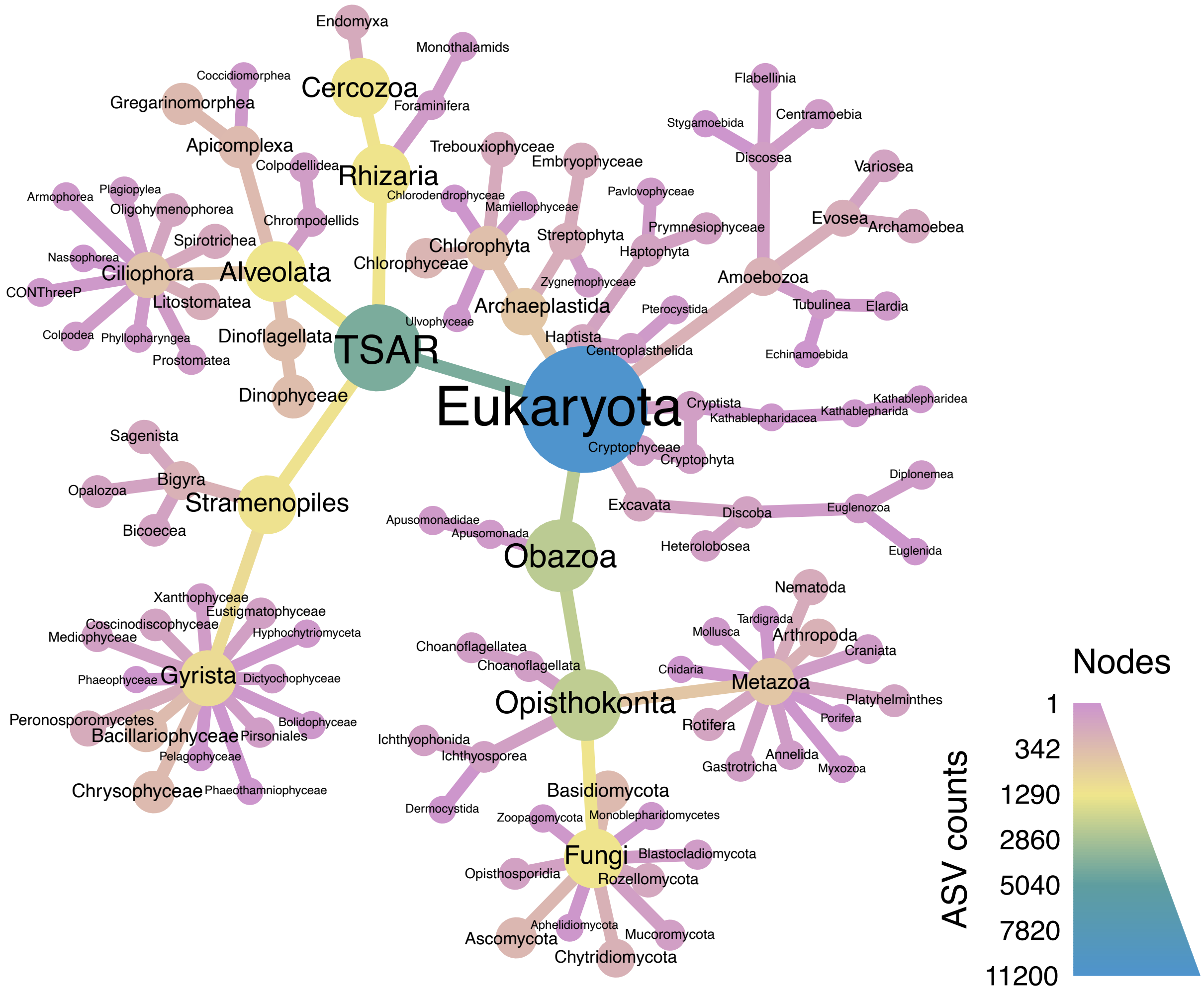


Supporting Information Fig. S6 Heattree showing the taxonomic diversity across all sediment samples from all lake cores.

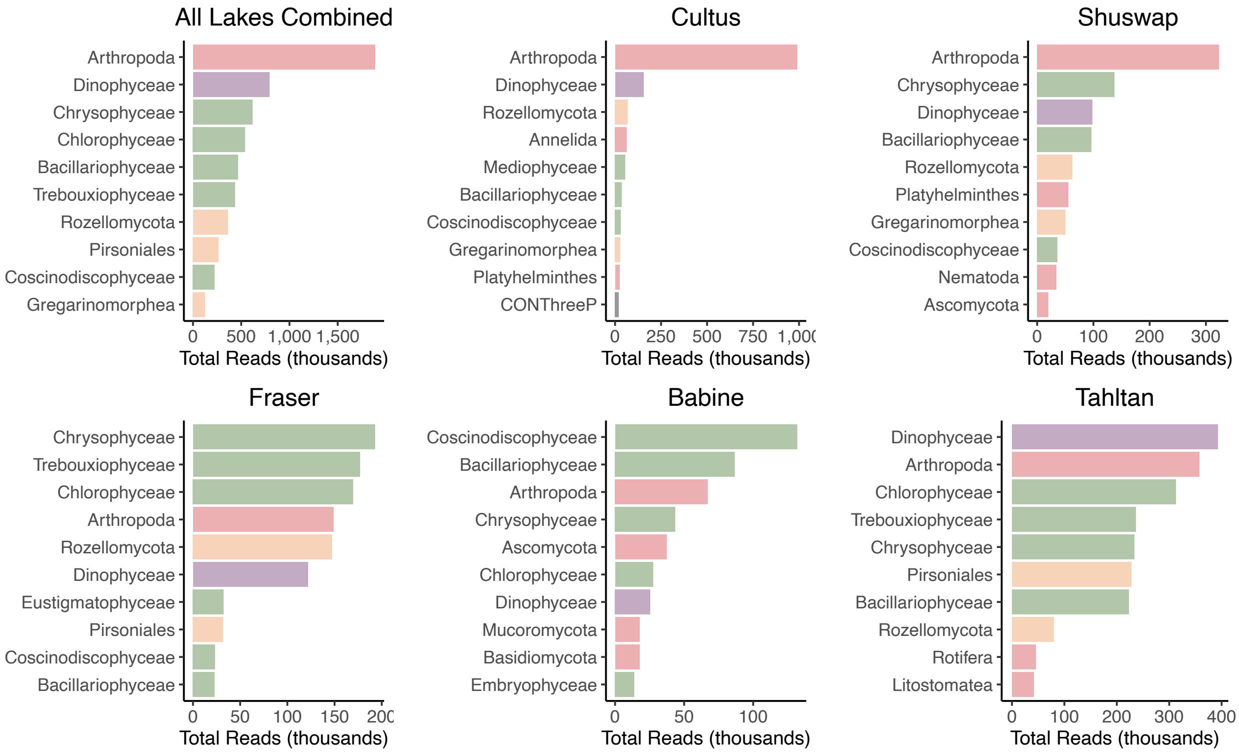


Supporting Information Fig. S7 Distribution of sequence reads at the class level.

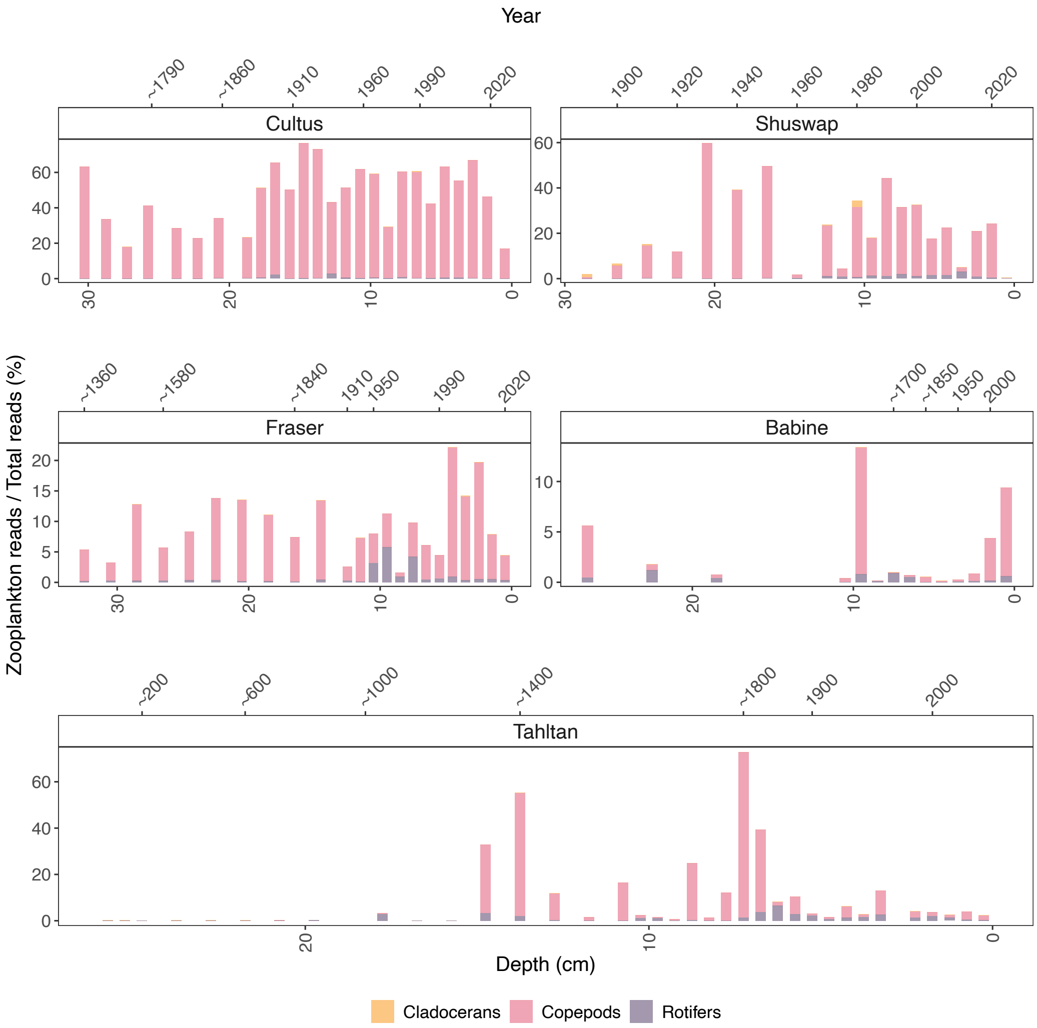


Supporting Information Fig. S8 Zooplankton bar chart showing the read proportions of three major zooplankton groups: cladocerans, copepods, and rotifers. Copepod communities were dominated by *Epischura* in Cultus and Shuswap (>99%), mixed composition in Fraser (78% *Epischura*, 22% *Leptodiaptomus*) and Babine (24% *Epischura*, 45% *Leptodiaptomus*), and dominated by *Leptodiaptomus* in Tahltan (>99%).

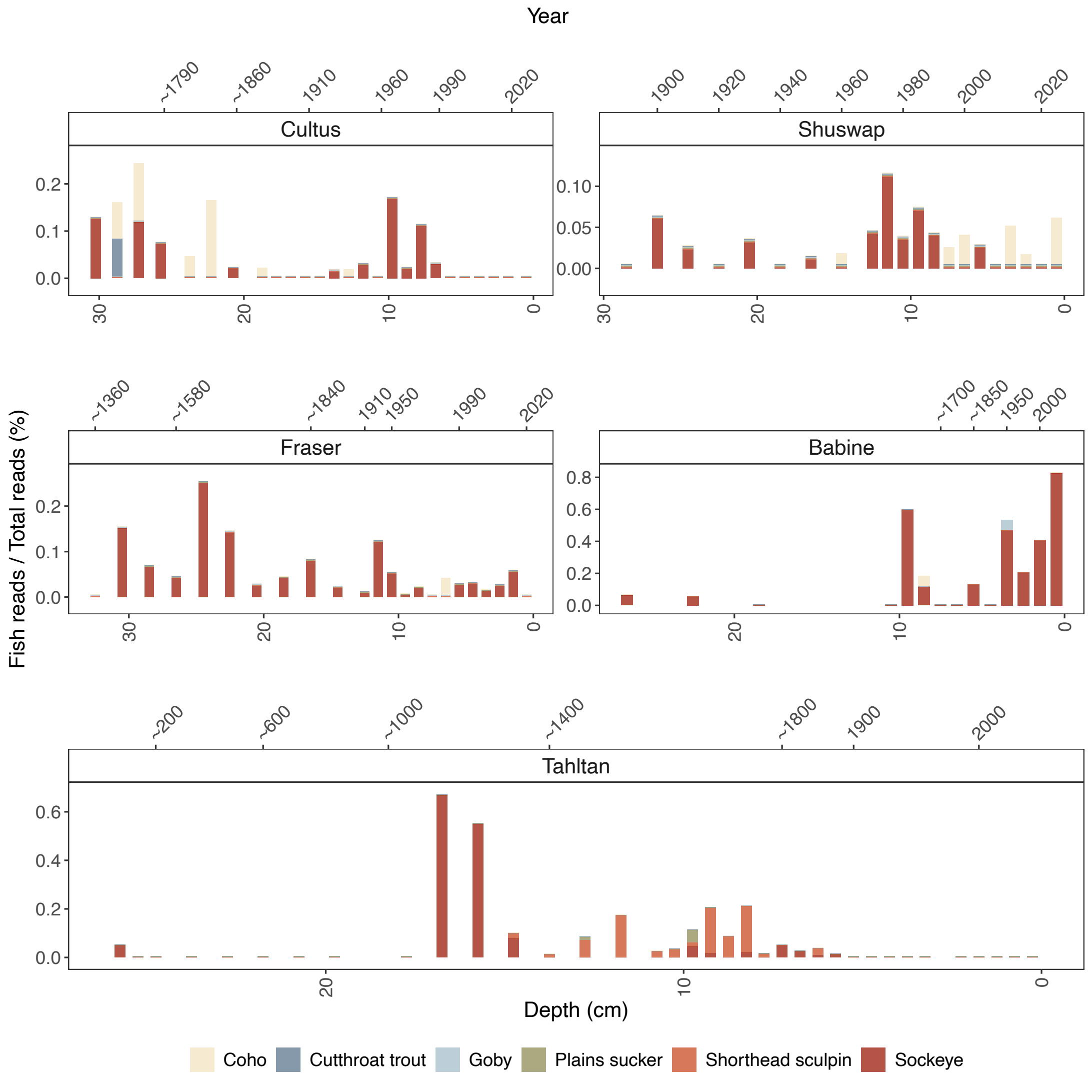


Supporting Information Fig. S9 Fish bar chart showing the read proportions of varying fish species.

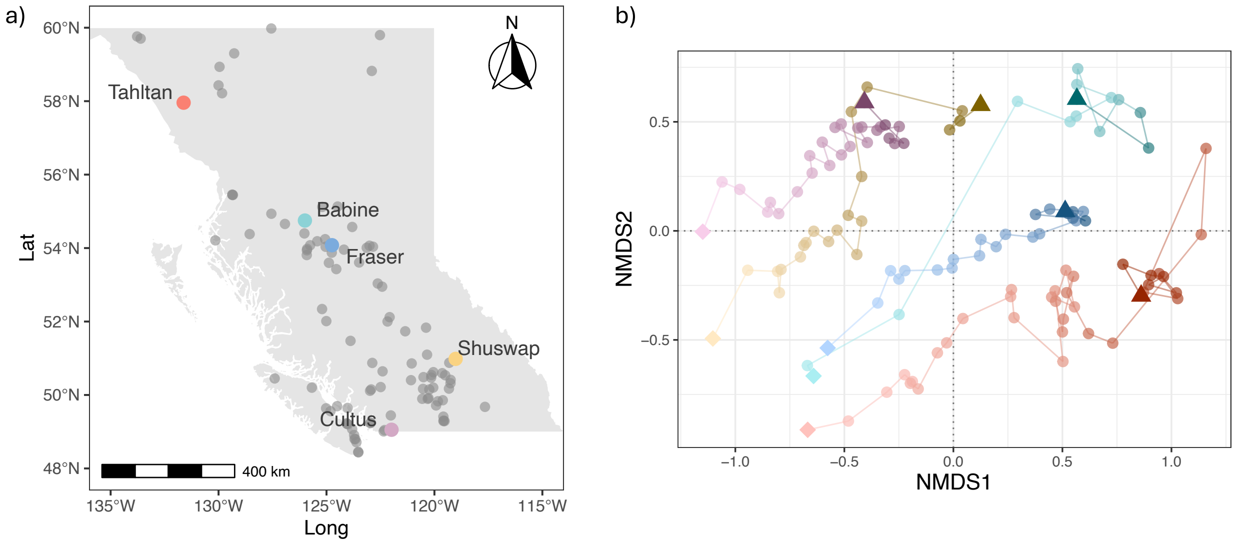


Supporting Information Fig. S10 LakePulse BC sampling sites (grey dots, *n* = 98) and Ordination of salmon nursery lakes retaining only those matched with the LakePulse dataset, with the surface sediments marked by a diamond and bottom sediments indicated by a triangle.

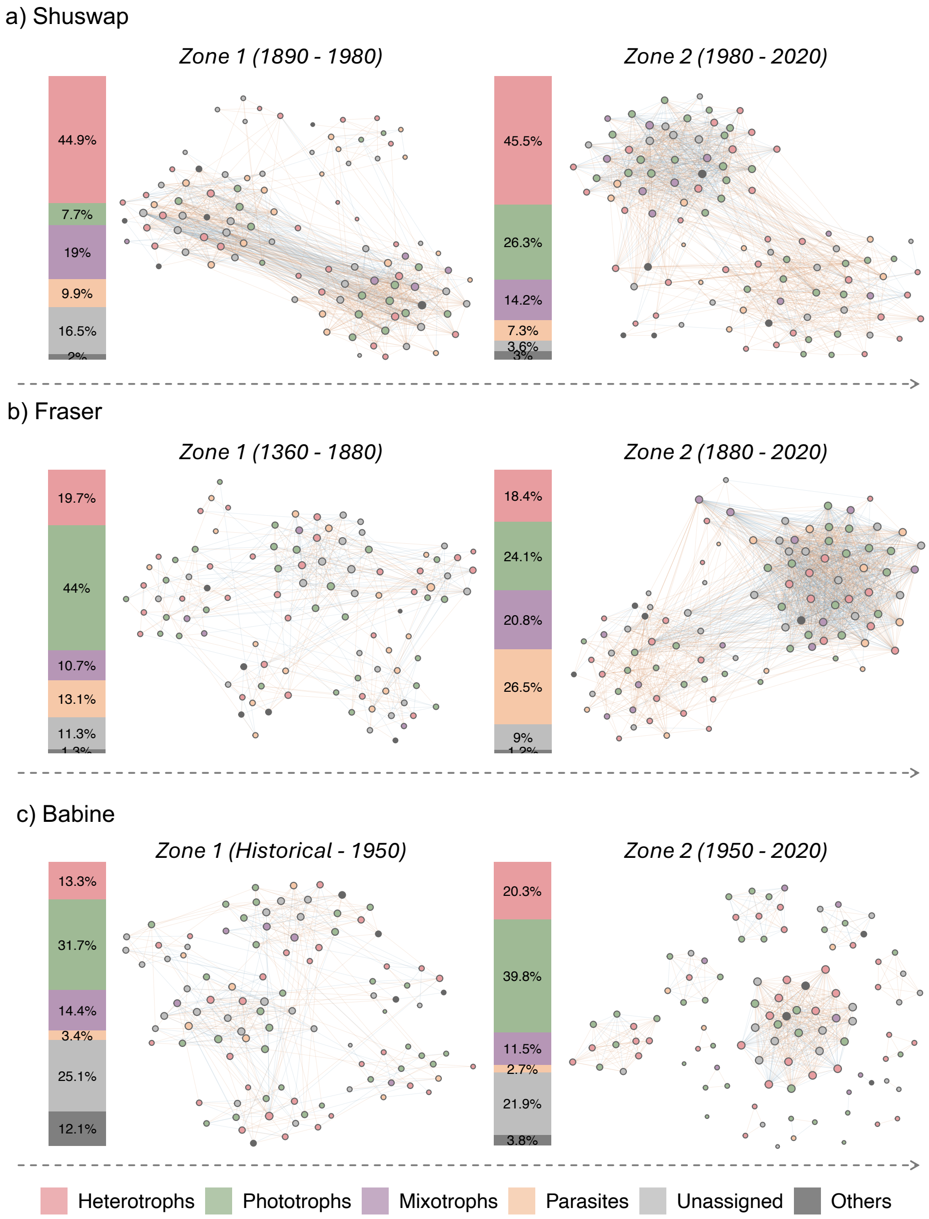


Supporting Information Fig. S11 Network analysis showing co-occurrence patterns of dominant ASVs in Shuswap, Fraser, and Babine lakes across different time zones. Red edges indicate positive correlations while blue edges represent negative associations. Bar plots show the relative proportions of dominant amplicons within each zone, classified into five trophic groups.

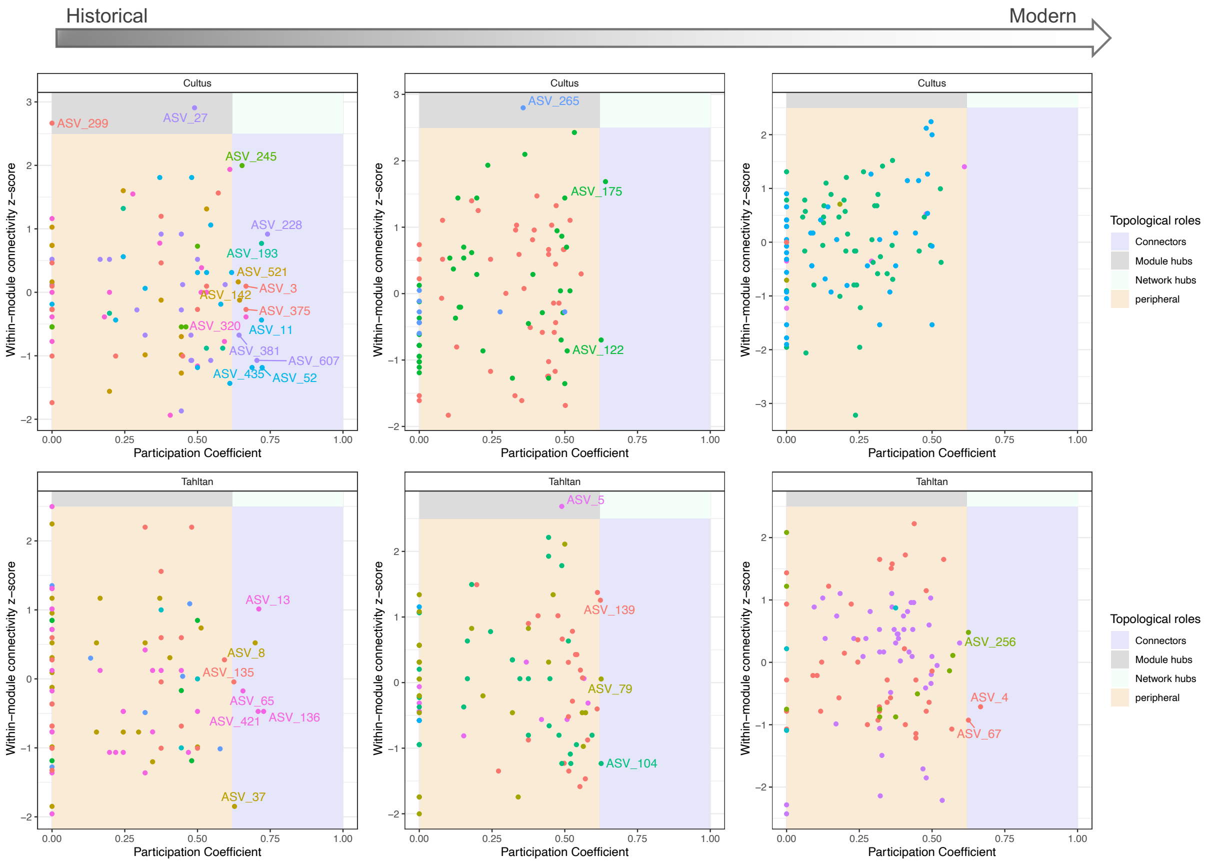


Supporting Information Fig. S12 Topological roles of each network for Cultus and Tahltan lakes. All ASVs that were identified with the topological roles of connectors, module hubs, or network hubs have been labeled.

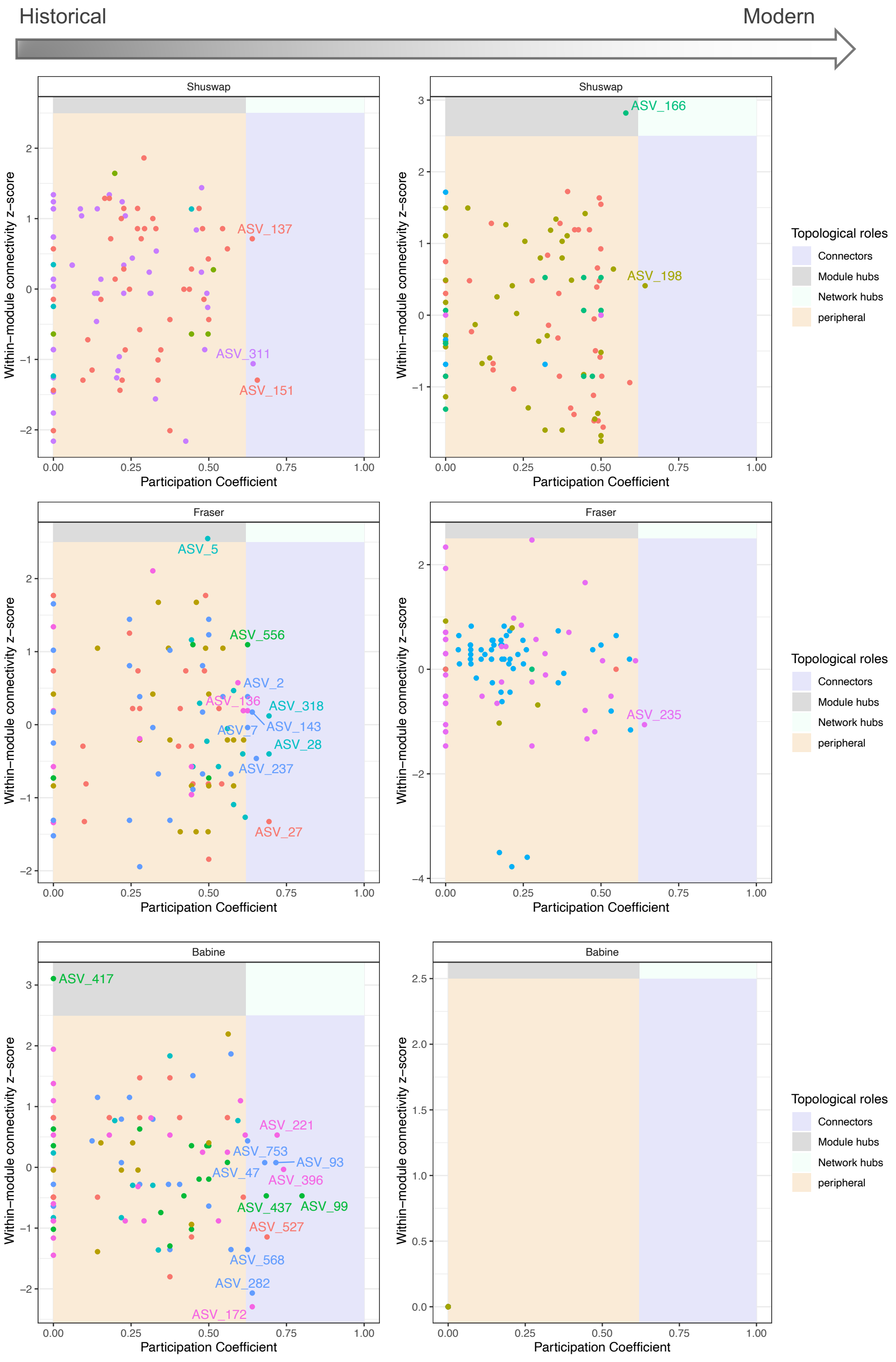


Supporting Information Fig. S13 Topological roles of each network for Shuswap, Fraser, and Babine lakes.

### Supporting Information S2. Lake-specific history

Here, we synthesized evidence from previous analyses to reconstruct the limnological histories of Cultus and Tahltan Lakes, representing the southernmost and northernmost systems in our study. Taxa with high ordination loadings were selected to guide ecological interpretations (Fig. S14), which are interpreted alongside historical records of human activity.

*Cultus Lake*

The ordination analysis revealed an ecological transformation from low nutrient, minimal human impact conditions toward intensive anthropogenic influences (Fig. 5b). A spike in Rhabdocoela (Fig. 8a) around the 1850s likely reflected increased organic matter from early watershed development, enhancing food availability for these predatory flatworms (Schockaert et al. 2008). During this period, the network remained relatively modular (Fig. 6a), suggesting ecosystem stability.

By the 1920s, extensive logging elevated organic carbon loading, shifting the lake toward the high-DOC conditions by mid-century (Fig. 5b). The zooplankton:phytoplankton ratio peaked during this period (Fig. 8a), potentially driven by elevated nutrient availability supporting zooplankton proliferation and/or decreased fish predation (Carpenter et al. 1985). Network modularity declined with increased positive co-occurrences (Fig. 6a), suggesting that under elevated stress, tolerant taxa persisted through cooperative rather than competitive dynamics.

The most recent era points to some improved conditions but new stressors as well. A pronounced *Tubifex* peak in the 1970s, comprising over 40% of reads (Fig. 8a), reflected polluted conditions (Aston 1973). Installation of a sewer system in 1979 likely contributed to the subsequent decline in sludge worms. By the 1990s, recreational pressure and climate change intensified, shifting the lake toward high human impact (HI) condition (Fig. 5b). Planktonic diatom abundance (especially *Cyclotella* sensu lato) increased markedly over the past two decades (Fig. 2b, 8a), likely reflecting altered winter-spring mixing rather than summer stratification dynamics, as monitoring data show dinoflagellates and chrysophytes dominate during stratification due to their mixotrophic abilities (Fig. 2a, 7b; Stoecker et al. 2017).

*Tahltan Lake*

Tahltan Lake exhibited a complex ecological trajectory over the past thousand years. Its initial high-nutrient position (Fig. 5f) likely reflects historically high salmon-derived nutrients, with the network showing high modularity suggesting stable niche partitioning (Fig. 6b).

Around ~1100 CE, a sharp C:N spike and sediment color shift from dark to light grey signaled major ecological transition (Fig. 8b). Elevated C:N suggests increased terrestrial organic matter input, while sediment color change indicates climatic/hydrological shifts. A concurrent *Gregarinomorphea* (invertebrate parasite) peak may reflect environmental stress during this disturbance. The ordination shows a shift toward higher ion concentrations (Fig. 5e), supporting altered hydrology. The transition back to organic-rich sediments is marked by a peak in the zooplankton:phytoplankton ratio in the early 19th century (Fig. 8b), suggesting recovery and restructuring of the aquatic community.

Pirsoniales (*Noirmoutieria* clade), parasitic protists preferring low-oxygen environments (Prokina et al. 2024), were historically dominant but declined with lower organic sediment content (Fig. 8b). They resurged between ~1700-1900 CE when organic-rich layers returned, and the lake shifted back toward its historical high-nutrient position (Fig. 5f), possibly linked to increased fish biomass. The most recent transition began around 1900 CE, when Pirsoniales declined coinciding with rising diatom sedDNA (Fig. 8b). Over the past century, the lake shifted into the left ordination space (Fig. 5f), likely reflecting longer growing seasons.

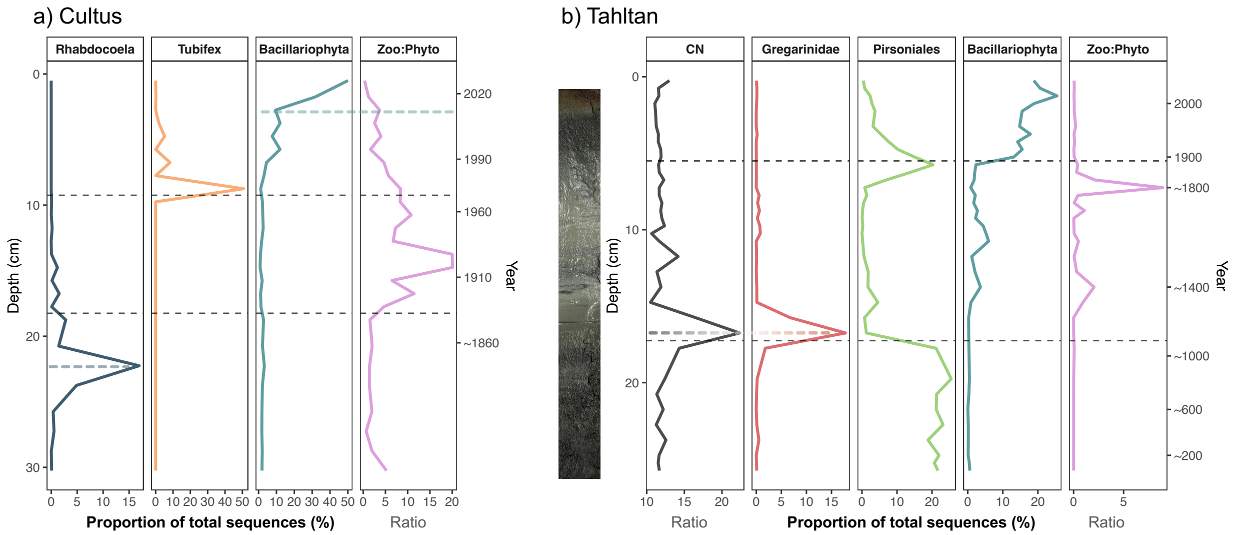


Supporting Information Fig. S14 Lake-specific stratigraphies highlighting major natural and human-induced environmental changes in a) Cultus and b) Tahltan Lakes. Black dash lines mark divisions of the major time zones. The following diagnostic organisms are highlighted: Rhabdocoela is an order of flatworms; *Tubifex* is a genus of annelids; Gregarinomorphea is a class of invertebrate parasites; Pirsoniales is an order of parasitic protists.

**Supporting Information References**


Aston, R. J. 1973. Tubificids and water quality: a review. Environmental Pollution 5: 1-10.

Balint, M. and others 2018. Environmental DNA Time Series in Ecology. *Trends in Ecology & Evolution* 33: 945-957.

Carpenter, S. R., J. F. Kitchell, and J. R. Hodgson. 1985. Cascading trophic interactions and lake productivity. *BioScience* 35: 634-639.

Prokina, K. I., N. Yubuki, D. V. Tikhonenkov, M. C. Ciobanu, P. Lopez-Garcia, and D. Moreira. 2024. Refurbishing the marine parasitoid order Pirsoniales with newly (re)described marine and freshwater free-living predators. *Journal of Eukaryotic Microbiology* 71: e13061.

Schockaert, E. R., M. Hooge, R. Sluys, S. Schilling, S. Tyler, and T. Artois. 2007. Global diversity of free living flatworms (Platyhelminthes, “Turbellaria”) in freshwater. *Hydrobiologia* 595: 41-48.

Selbie, D. T. 2008. Low-frequency climatic and solar forcing of Northeast Pacific salmon production, p. 163-186. In D. T. Selbie, *Large-scale exogenous forcing of long-term Pacific salmon production and ecosystem interactions in western North America*. Ph.D. thesis. Queen's University.
